# Mapping molecular datasets back to the brain regions they are extracted from: Remembering the native countries of hypothalamic expatriates and refugees

**DOI:** 10.1101/307652

**Authors:** Arshad M. Khan, Alice H. Grant, Anais Martinez, Gully A.P.C. Burns, Brendan S. Thatcher, Vishwanath T. Anekonda, Benjamin W. Thompson, Zachary S. Roberts, Daniel H. Moralejo, James E. Blevins

## Abstract

This article, which includes novel unpublished data along with commentary and analysis, focuses on approaches to link transcriptomic, proteomic, and peptidomic datasets mined from brain tissue to the original locations within the brain that they are derived from using digital atlas mapping techniques. We use, as an example, the transcriptomic, proteomic and peptidomic analyses conducted in the mammalian hypothalamus. Following a brief historical overview, we highlight studies that have mined biochemical and molecular information from the hypothalamus and then lay out a strategy for how these data can be linked spatially to the mapped locations in a canonical brain atlas where the data come from, thereby allowing researchers to integrate these data with other datasets across multiple scales. A key methodology that enables atlas-based mapping of extracted datasets – laser-capture microdissection – is discussed in detail, with a view of how this technology is a bridge between systems biology and systems neuroscience.

## 1: Introduction

### 1.1: Summary and rationale

In this article, we envision ways in which molecular information extracted from the brain using methods such as transcriptomics, proteomics, and peptidomics can be anchored to locations in standardized atlas maps of the brain in order to preserve the provenance of the datasets and contextualize them with other datasets. We argue that whereas most researchers probe, dissect, mine, or interrogate the living brain and report back with valuable scientific information, such information would be worth more if it included *mapped locations* of where they traveled and what they found there. Mapping to a standardized reference allows current and future travelers to return to the same landscape with accuracy and precision, generate reproducible data from reproducible experiments, and allows them further to integrate and contextualize new data they gathered in that mapped location with other data gathered in the same space. By carefully documenting the locations, for example, of brain regions from which molecular information is extracted for large-scale analyses, scientists can contribute further to our collective history of the native landscape from which this expatriated molecular information originated.

### 1.2: Topic and organization

We have chosen to use the hypothalamus as an exemplar structure to illustrate the possibilities of such an effort, a choice that is predicated in part on our own experiences in mapping and modeling multi-scale data for this brain region (e.g., Khan et al. 2006; Khan, 2013; Zséli et al. 2016; Khan et al. 2017; 2018), and because a review of “-omics” work on the hypothalamus in the context of spatial mapping has not yet, to our knowledge, been attempted. So far, molecule extraction from hypothalamus has been focused primarily on mining either the whole hypothalamus or its well-defined sub-regions to the virtual exclusion of parts that are less well understood. If wider and more systematic sampling of areas within the hypothalamus were to be conducted, atlas mapping efforts will play an even greater role in helping us understand the organization of those areas that remain poorly defined. The additional benefit of mapping molecular data to a standardized atlas is that the data can be contextualized with multi-scale datasets mapped to the same reference map.

Below, following a brief exploration of the biological importance of location information in the brain (**Section 2**), we summarize the historical antecedents to current molecular extraction work done on the brain (**Section 3**) and the hypothalamus specifically (**Section 4.1**), focusing on those datasets that include spatial data about the regions extracted. We then survey studies that have examined the molecular landscape of the hypothalamus using transcriptomics, proteomics and peptidomics (**Section 4.2**). The rationale behind the separation of proteomics from its subdomain, peptidomics, is based on the fact that the latter involves analytical procedures that are distinct from those in general proteomics, including more rigorous purification and more comprehensive identification procedures (Schrader et al. 2014; Romanova and Sweedler 2015). The differences are great enough in methodology and concept that a separate consideration of peptidomic studies is warranted. The narrative then shifts to specific strategies that we envision will be required, especially the technique of laser-capture microdissection (LCM) (**Section 5**), to enable the accurate mapping of hypothalamic molecular datasets to a standardized atlas of the brain (**Section 6**), and the benefits of such mapping (**Section 7**). We conclude with a view to current and future directions for this research (**Section 8**).

## 2: Why does location matter?

The brain is a very heterogeneous organ that contains diverse, non-repeating, and non-redundant sub-regions (e.g., see Balázs et al. 1972; Lehrer and Maker 1972; Palay and Chan-Palay 1972). Studies in many animal model systems have now revealed that brain region is a major determinant of gene expression patterns. Therefore, the location of areas sampled using “-omics” technologies will determine critically the complement of molecules expressed. Left- and right-handedness in cichlid fish, for example, is correlated strongly with hemispheric and regional asymmetry of gene expression (H. Lee et al. 2017). In songbirds, clustering analyses performed on retrieved sets of genes demonstrate a strong association of gene expression with brain region (Replogle et al. 2008; Drnevich et al. 2012; Balakrishnan et al. 2014). This also holds true for mammalian brain. Even between strains of mice (which can exhibit size differences for the whole brain and for individual brain regions: Badea et al. 2009), one report has estimated a 1% difference in baseline expression patterns in at least one brain region, and that gene expression differences in response to a physiological perturbation (in this case, seizure) produce marked differences in gene expression patterns in brain regions between strains (Sandberg et al. 2000). A re-analysis of the datasets of Sandberg et al. (2000) by Pavlidis and Noble (2001) reveals even greater differences in regional variation among the genes between the strains. These observations were extended by Nadler et al. (2006), who found, across ten inbred mouse strains, that there was a nearly 30% difference in gene expression in at least one brain region among those examined. Robust strain differences have also been documented for transcripts enriched in the rat hypothalamic neurohypophysial system (Hindmarch et al. 2007). Moreover, Dong et al. (2009) show that specific patterns of gene expression are associated with specific domains where distinct neural projection patterns emerge within the hippocampus, and Wolf et al. (2011) show that there is a strong predictive association of neural connections and gene expression within specific brain regions (also see Sun et al. 2012). Superimposed on this complexity are strain-dependent variations in the sexual dimorphism of certain brain nuclei (Robinson et al., 1985; Mathieson et al. 2000), and differences in how gene expression networks in the brain are modulated as a result of expression quantitative trait loci (eQTLs) that are sex-specific (Mozhui et al. 2012; also see Pandey and Williams, 2014). Thus, it is important to consider just what we as scientists lose if we endeavor to extract molecular information from the brain without attempting to preserve the provenance of where the extraction took place. Before addressing this issue more directly, it is useful to survey the history behind efforts to identify chemical and molecular information encoded in the brain.

## 3: Historical antecedents

### 3.1: Heuristic entry points to relevant history

Recent “-omics” work has been informed to various extents by seminal works conducted during the last 150 years which we have categorized heuristically along major research themes: *composition*, *communication*, *reaction* and *localization*. First, regarding *composition*, our current effort to understand dynamic changes in the expression of genes and proteins in the nervous system is predated by work that first identified its fundamental chemical (elemental) constituents (e.g., Thudicum 1884; Richter 1957; McIlwain 1959; Friede 1966). Studies of the molecular constituents of neural machinery were motivated in part by contemporaneous questions concerning the ionic and chemical bases of muscle and nerve excitability (Helmholtz 1850; Nernst 1888; Overton 1902a,b; Hill 1932; Hodgkin and Huxley 1939; 1945; 1952a–f; Hodgkin et al. 1952; Fatt and Katz 1952; see various reviews by Boring 1942; Kleinzeller 1999; Häusser 2000; Bennett 2001; Huxley 2002; De Palma and Pareti 2011; Schwiening 2012; also see Khan 2009). Predating current work on proteomics and peptidomics, work on chemical composition was also marked by efforts in the 1980s by Tatemoto and colleagues to use chemical methods to isolate, identify and determine the sequence of neuropeptides such as galanin and neuropeptide Y (Tatemoto and Mutt 1980; Tatemoto et al. 1982; 1983; see Schrader et al. 2014).

Second, concerning *communication*, the mining of molecules coding for neurotransmitter and neuropeptide machinery in the nervous system finds its antecedents in both Bayliss and Starling’s (1902) discovery of peptide hormone secretion from the pancreas (also see Hirst 2004; Schrader et al. 2014), and Loewi’s (1921) discovery of cholinergic neurotransmission in the peripheral nervous system. Ensuing efforts to gather evidence for a role for acetylcholine as a neurotransmitter in the central nervous system (e.g., Feldberg 1952) were facilitated by histochemical methods (Koelle and Friedenwald 1949; Anglade and Larabi–Godinot 2010; but see Levey et al. 1983), which helped contribute to the maturation of chemical neuroanatomy as a subdiscipline of neuroanatomy (also see: Jacobowitz and Palkovits 1974; Palkovits and Jacobowitz 1974). Importantly, histochemistry became useful to trace metabolic turnover in the brain, since it was performed on living tissue and was based on enzymatic activities catalyzing the conversion of substrates to detectable products.

This work complemented contemporaneous studies – grouped thematically under *reaction* – that concerned the metabolism of living neural tissue, pioneered by Warburg, McIlwain and others (e.g., see Warburg et al. 1924). Finally, a fourth long-standing body of work that informs “-omics” approaches concerns the historical quest to understand how various functions of the brain are derived from specific locations within its complex structure, the theme of *localization* (e.g., Flourens 1824; Ferrier 1873; Adrian 1939; also see Swanson 2007). This theme directly informs efforts to isolate portions of the nervous system for detailed study through careful extraction and sampling, a topic we delve into next.

### 3.2: Sampling at the level of the single cell

Early interests in sampling very small portions of the central nervous system prefigure current interests in developing “spatially resolved” approaches (e.g., Crosetto et al. 2015) for transcriptomics and proteomics of neural tissues. Otto Deiters (1865) famously provided anatomical descriptions emphasizing the emergence of a single axon and multiple dendrites from motor neuron cell bodies in the spinal cord (also see Deiters and Guillery 2013), which he isolated individually by hand from chromic acid- or potassium dichromate-hardened (i.e., fixed) tissue (**Fig. 1A**). Several investigators such as Hans Held and others followed suit using a variety of fixed preparations to study neurons in greater detail (see introductory comments in Chu (1954) for an overview).

**Fig. 1.**
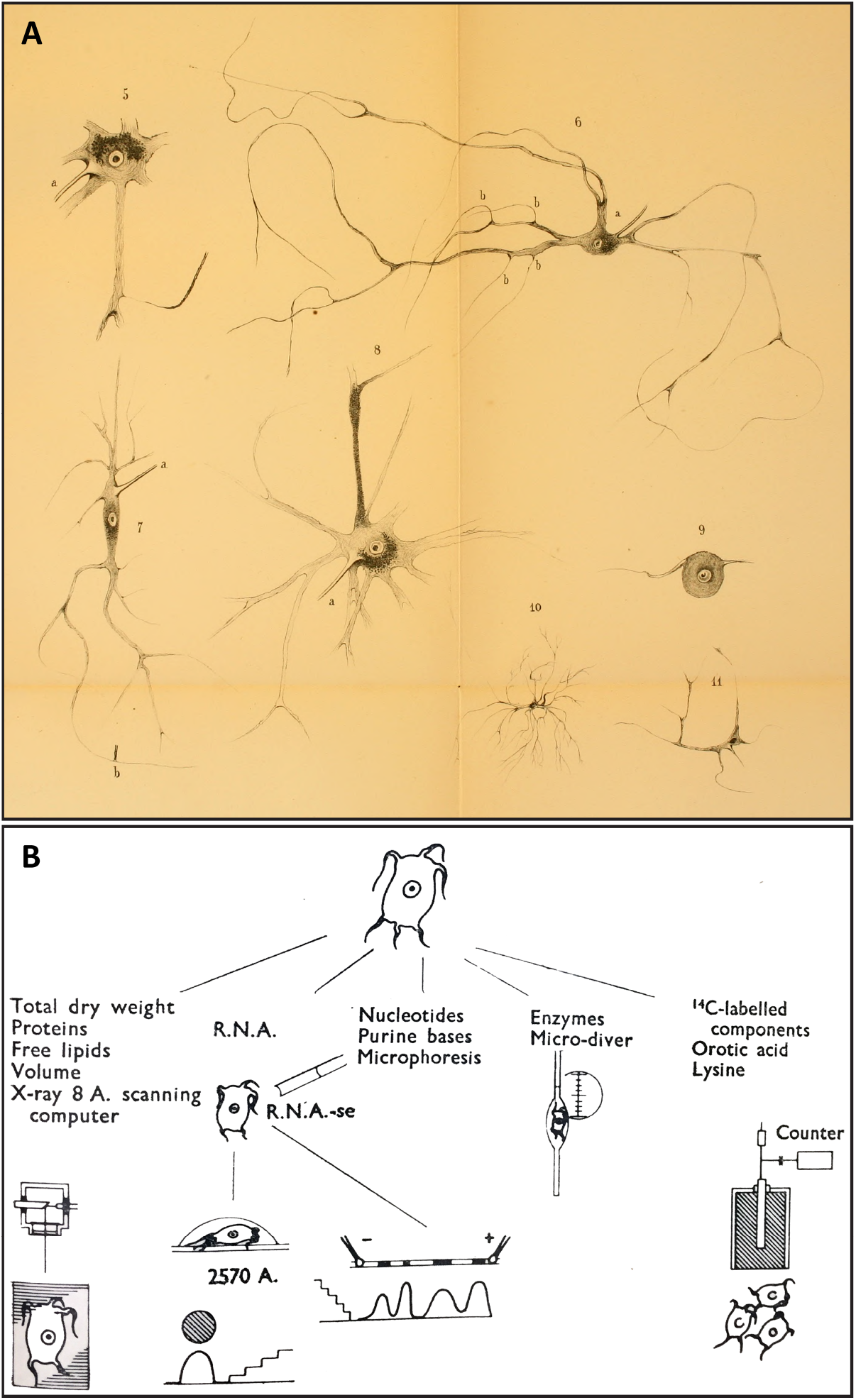
Demonstrations of single-neuron isolation procedures, nearly a century apart, which anticipate isolation methods performed at present for studying single-cell transcriptomics, proteomics, and peptidomics. **(A)**. *Plate II* of Deiters (1865), showing the morphologies of neurons collected from the central nervous system by hand microdissection. Note the recovery of many fine processes extended from each perikaryon, including axons and dendrites. These drawings are in the public domain. **(B)**. *Figure 2* of Hydén (1959) showing a workflow schematic of possible assays that can be performed on single neurons isolated from freshly prepared tissue of the lateral vestibular nucleus. Neurons in this region are named “giant cells of Deiters” in honor of Deiters’s initial description of these cells (see Sotelo & Palay, 1968). Reproduced with permission from Nature Publishing Group.

Deiters’ manual single-cell microdissection technique anticipates, by almost a century, single cell isolation from *fresh* neural tissue preparations pioneered by Ezio Giacobini (1956) for frog, rat and cat spinal cord and peripheral ganglia; and Holger Hydén (1959) for the mammalian brain. Hydén, for example, used manual microdissection to isolate and chemically analyze (fittingly) the “giant neurons of Deiters” found in the lateral vestibular nucleus (**Fig. 1B**; Hydén 1959; also see Hydén 1967; Sotelo and Palay 1968; Rose 1999). Along with Giacobini’s and Hydén’s work using freshly microdissected neurons, related methods developed by Lowry (1953), Chu (1954), Roots and Johnston (1965), Johnston and Roots (1966) and others using fixed, freeze-dried, and reagent-impregnated tissues ushered in an era of “micro-chemical methods”, in which a variety of *chemical assays* could be performed on single cells isolated from various regions of the central nervous system (cogently reviewed by Johnston & Roots, 1970; also see Roberts and Baxter 1963; Osborne 1974). Eberwine et al. (1992) performed gene expression analysis on individual, freshly dissociated hippocampal neurons. More recently, single-cell isolation has now been conducted using laser-capture microdissection (LCM) methods (e.g., D. Williams et al. 2008; Blevins et al. 2009; Briski et al. 2010; Carreño et al. 2011; Landmann et al. 2012) or cell sorting methods (e.g., Draper et al. 2010; Henry et al. 2015; Macosko et al. 2015; R. Chen et al. 2017; S. Chung et al. 2017; Mickelsen et al. 2017; Romanov et al. 2017; also see Okaty et al. 2011; Poulin et al. 2016). Thus, single cell isolation methods first used for the purposes of morphological and structural investigation evolved for use in biochemical, molecular and functional analyses.

### 3.3: Sampling at the level of isolated tissues

Alongside single-cell isolation methods were those procedures driven by the need to examine metabolically active states of the nervous system in isolated tissue preparations where the local microenvironment of the cells was, to some extent, still maintained. Metabolic studies of living tissues maintained in isolation were pioneered by Otto Warburg’s laboratory in the 1920s, including studies performed on the isolated retina (Warburg et al. 1924).

## 4: Molecular mining of the hypothalamus

### 4.1: Early studies

Prior to the advent of high throughput methods, several laboratories performed a variety of techniques to isolate and examine the molecular constituents of the hypothalamus, either using living samples or fixed samples *post mortem*. A number of such studies were conducted because investigators at the time were motivated to differentiate the functions of the pituitary gland from the overlying hypothalamus (e.g., see Lisser 1927). Other investigators concerned themselves more with trying to understand, through histochemistry, the nature of chemical transmission in the hypothalamus (reviewed by Pilgrim 1974), to validate, for example, the existence of cholinergic neurotransmission within hypothalamic regions (*see* **Section 2.1**). Feldberg and Vogt (1948) isolated the supraoptic hypothalamic nucleus in the dog to perform acetylcholinesterase (AchE) histochemistry, a method also performed in hypothalamus by Abrahams et al. (1957). Still others extended the tradition of Warburg and colleagues by examining the living hypothalamus for insights into metabolic processes occurring within this tissue, primarily through the use of radiolabeled phosphate incorporation. For example, Borell and Örström (1945) examined radiolabeled phosphate accumulation in the anterior and posterior portions of the hypothalamus, and Roberts and Keller (1953, 1955) studied glycolysis in hypothalamic tissue preparations. Bakay (1952) examined radiolabeled phosphate incorporation in the human hypothalamus *post mortem* following the deaths of terminally ill cancer patients who had received intravenous tracer to track their brain tumors.

In what is perhaps the earliest demonstration of chemical analysis performed on an explicitly defined microdissected sub-region of the hypothalamus, Forssburg and Larsson (1954) sampled a portion of the hypothalamus from male and female rats that were either food-deprived for 24 h or *ad libitum-fed* and that received radioactive (^14^C; Na_2_H^32^PO_4_) tracer injections to track their carbon and phosphate metabolism. Brains were rapidly dissected and frozen, and 20–50 μm-thick sections were obtained of the brain, and examined carefully for the incorporation of ^14^C and ^32^P in chemically extracted fractions of the microdissected tissue. Importantly, the authors included a schematic to outline the areas they micropunched (**Fig. 2A**), including areas they sampled outside of the hypothalamus that served as a control. Their careful documentation of the sampled area and use of a custom-made micropunch tool (which they also illustrated in their study) anticipates the later use of similar instruments as developed by Palkovits and colleagues to sample discrete parts of the brain (see Palkovits 1973; 1975; 1986; 1989; Jacobowitz 1974; also see Jacobowitz 2006).

**Fig. 2.**
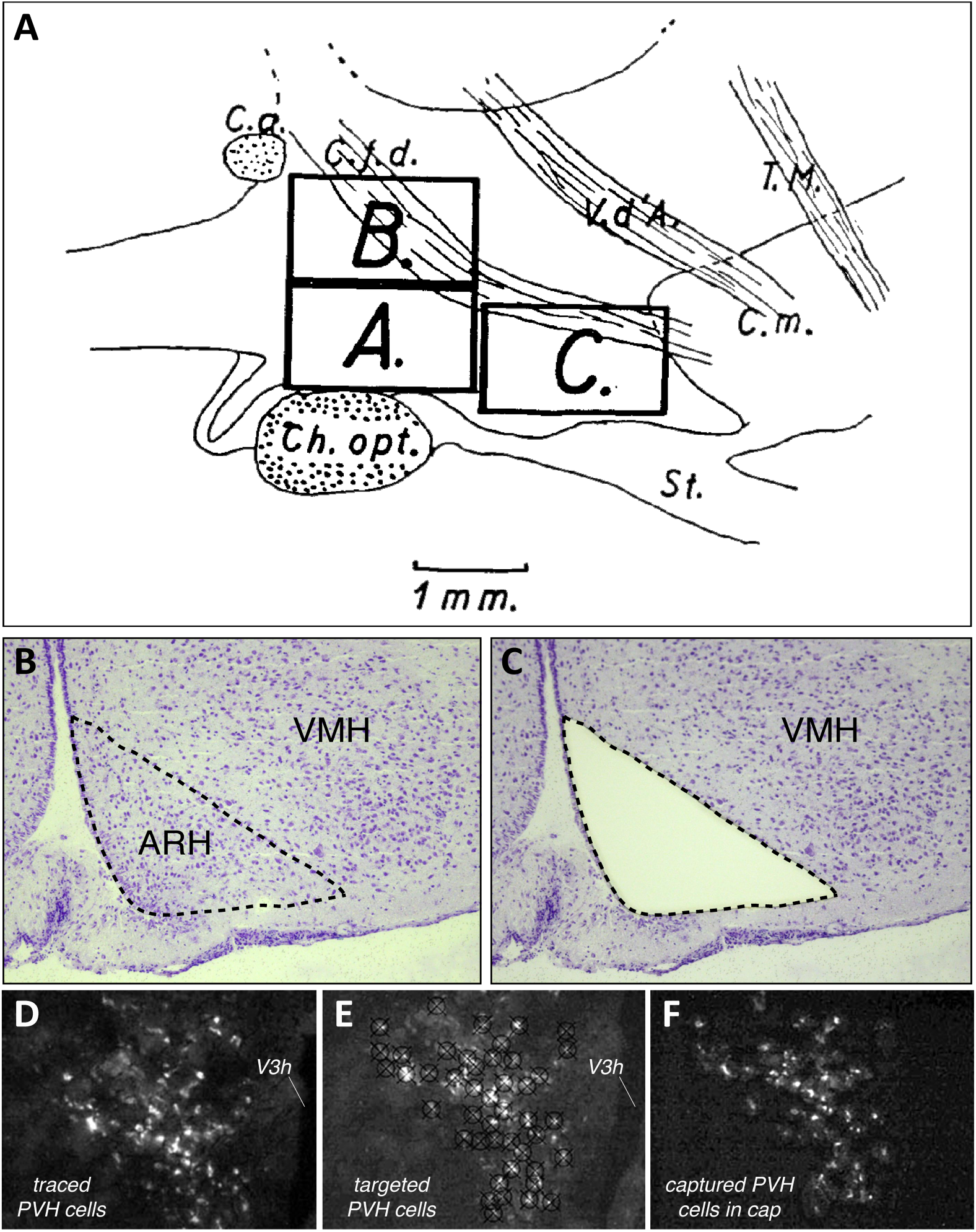
Examples of documentation, past and more recent, of microdissected areas sampled from rat hypothalamus for chemical or molecular analyses – from gross micropunch (**A**), to region-level laser-capture microdissection (LCM) **(B, C)**, to single-cell LCM **(D–F)**. **(A)**. *Figure 14* of Forssburg and Larsson (1954), showing a sagittal drawing of the hypothalamus, as defined by major fiber tract landmarks: Ch. Opt. = optic chiasm; C.a. = anterior commissure; C.f.d. = fornix; V.d’A. (*Tract of Vicq d’Azyr*) = mammillothalamic tract; T.M. (*Tractus Meynert*) = fasciculus retroflexus; St. = infundibular stalk. The boxes denoted by letters mark the regions micropunched from thin frozen sections at the locations indicated by the drawing, with A and B serving as control regions and C as the region of interest containing the lateral and ventromedial hypothalamus. Reproduced with permission from John Wiley & Sons, Ltd. **(B, C)**. A Nissl-stained view of the arcuate hypothalamic nucleus (ARH) and ventromedial hypothalamic nucleus (VMH) in rat brain tissue sectioned in the coronal plane, before **(B)** and after **(C)** the tissue was subjected to LCM. The dotted outline marks the region captured by the LCM instrument; note how the Nissl pattern helps to delineate the boundaries of the region to be sampled, and the remaining tissue after LCM can then be used to map the sampled region to a digital atlas. These images are provided courtesy of Dr. Rebecca Hull and Nishi Gill (*see Acknowledgments*). **(D–F)**. Example of single-cell LCM of hypothalamic cells. These panels show photomicrographs adapted from *Figure 5* of Blevins et al. (2009), in which paraventricular hypothalamic (PVH) cells projecting to the hindbrain (as revealed in **(D)** by the presence of the retrograde tracer, cholera toxin subunit B (*white*), in PVH cells); have been targeted for LCM; see cross-patterns in **(E)**); and then have been collected into a microcentrifuge cap following LCM capture **(F)**.

Using these micropunch methods, and leveraging refinements (O’Farrell 1975) of the original two-dimensional gel electrophoresis method (Smithies and Poulik 1956) that allowed proteins to be separated by their apparent molecular weights and isoelectric points (reviewed by Dunn 1987), Jacobowitz and colleagues pioneered the systematic study of proteins from discrete micropunched regions of the rat brain, including from within the hypothalamus (Heydorn et al. 1983). Importantly, their study included a schematic of atlas maps from the rat brain atlas of König and Klippel (1963) to identify the approximate locations and diameters of their tissue micropunches. Among the many brain regions sampled were the anterior, paraventricular, ventromedial and dorsomedial hypothalamic nuclei. Although the authors were able to obtain apparent molecular weights, isoelectric points and relative amounts of proteins from their tissue punches (see also Heydorn et al. 1986), their study does not specifically identify the proteins themselves except in a few cases. Methods to do so, involving annotated databases, had not yet been developed. While micropunch methods continue to remain popular (e.g., see Atkins et al. 2011; Kasukawa et al. 2011), finer-grained studies that require more precise sampling of brain regions utilize LCM (Emmert-Buck et al. 1996), which is described in greater detail in **Section 5**, and a product of which is shown in Panels B and C of **Figure 2**. This higher resolution sampling using LCM has now been performed at the level of single hypothalamic cells (*e.g*., see **Figure 2D–F**).

### 4.2: Studies of the hypothalamus using high throughput methods

**Tables 1–3 (located after the References section)** summarize selected studies performed to extract molecular data from the hypothalamus using high-throughput transcriptomic, proteomic, and peptidomic approaches; respectively. Transcriptomic approaches include microarray (Fodor et al. 1991; Makoso and Southern 1992; also see Lenoir and Gianella 2006; Pirrung and Southern 2014) and next-generation sequencing (RNA-Seq; e.g., Mortazavi et al. 2008) technologies; proteomic and peptidomic approaches include protein separation methods such as electrophoresis and profiling technologies based on mass spectrometry (Gauss et al. 1999). A few of the tabulated studies are discussed below, beginning with studies which examined the hypothalamus as part of larger whole brain and/or multi-regional studies, and then on to studies in which the hypothalamus itself or its sub-regions were the main focus. Before these studies are examined in greater detail, it is useful to first consider the “state of the field” as a whole in terms of how much sampling of the hypothalamus and its various regions have been undertaken thus far. **Figure 3** is a snapshot of the level of coverage reported by the studies listed in **Tables 1–3**, organized by high throughput method and by spatial location within the hypothalamus. Specifically, a choropleth flatmap of the rodent brain, adapted from Swanson (2004), is utilized to highlight the degree to which either the whole hypothalamus (**Fig. 3A**), or individual sub-regions of the hypothalamus (**Fig. 3B–D**) have been sampled using transcriptomic, proteomic and peptidomic methods.

**Fig. 3.**
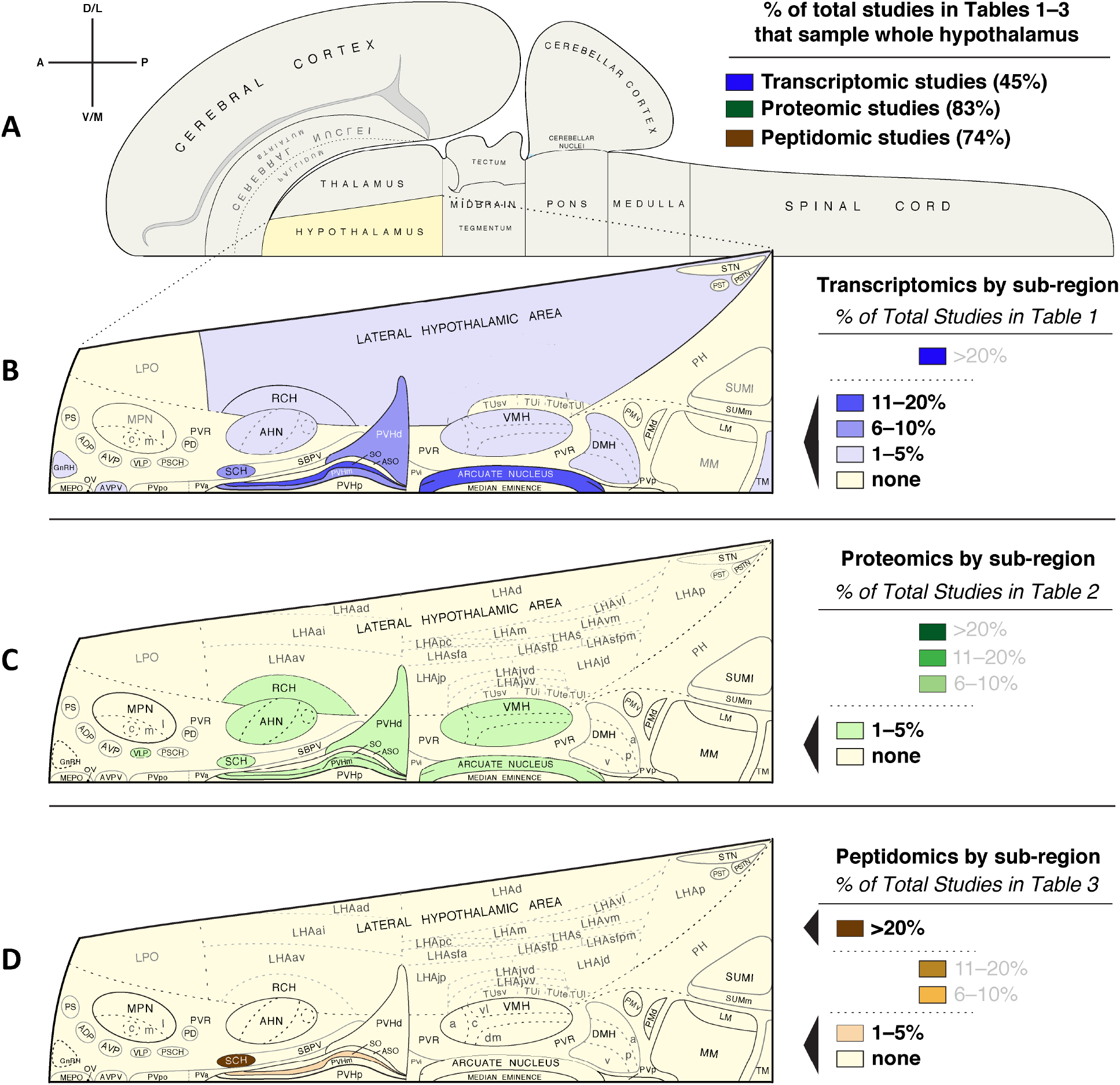
Survey of coverage for the hypothalamus or its various regions by published transcriptomic, proteomic and peptidomic studies listed in **Tables 1–3**. **(A)** Choropleth flatmap of the rat central nervous system (CNS), modified from Swanson (2004), illustrating the various major CNS subdivisions, including the hypothalamus. Note the *legend* at the upper left, which indicates directions of orientation (A, anterior; P, posterior; D/L, dorsal/lateral; V/M, ventral/medial). The *chart* to the right of the flatmap in **(A)** lists the % of total studies reported in each of the tables in this review (**Tables 1–3**) that conducted transcriptomic, proteomic, or peptidomic studies of the *whole* hypothalamus, respectively. **(B–D)** A breakdown, by *hypothalamic region*, of the percentage of studies in which a particular region was sampled for analysis, with choropleth flatmaps in **B** showing all hypothalamic regions analyzed by transcriptomic analyses, in **C** showing all proteomic studies, and in **D** all peptidomic studies. The percentage ranges are coded by colors and reflect percentages of the total number of studies listed in each table. Note that although rat brain flatmaps are used here, the studies are across many different taxonomic groups, including fish, chicken, goose, cow, pig, sheep, shrew, mouse, rat, guinea pig, hamster, dog and human. Therefore, the maps are meant to be convenient vehicles to convey a sense of the amount of coverage in the literature for any particular region, differences in their neuroanatomy or cytoarchitectural boundaries notwithstanding. Note also that for many shaded regions, the smaller abbreviations have been removed for sub-regions, to emphasize that studies did not sample at that level of resolution. Thus, the lateral hypothalamic area (LHA) may have been sampled, but the LHAjvv was not. Conversely, although large areas are shaded, in certain cases only a few cell types were specifically mined from the region rather than the region sampled as a whole, but this is not reflected in the diagrams. For explanation of all abbreviations, please see abbreviations list. The flatmaps from Swanson (2004) (and available at https://larrywswanson.com) are reproduced here under the conditions of a Creative Commons BY-NC 4.0 license (https://creativecommons.org/licenses/by-nc/4.0/legalcode).

**Table 1.**
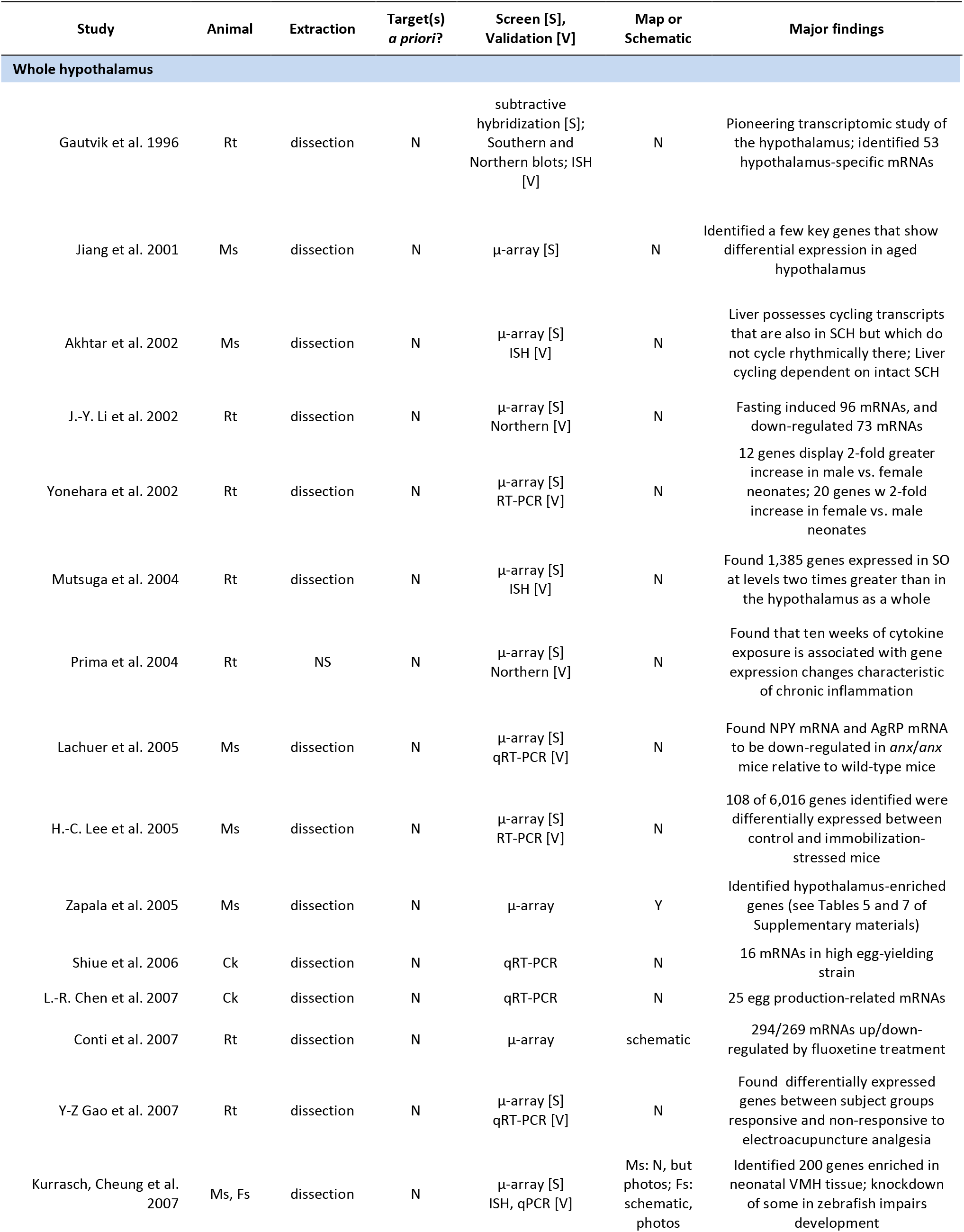
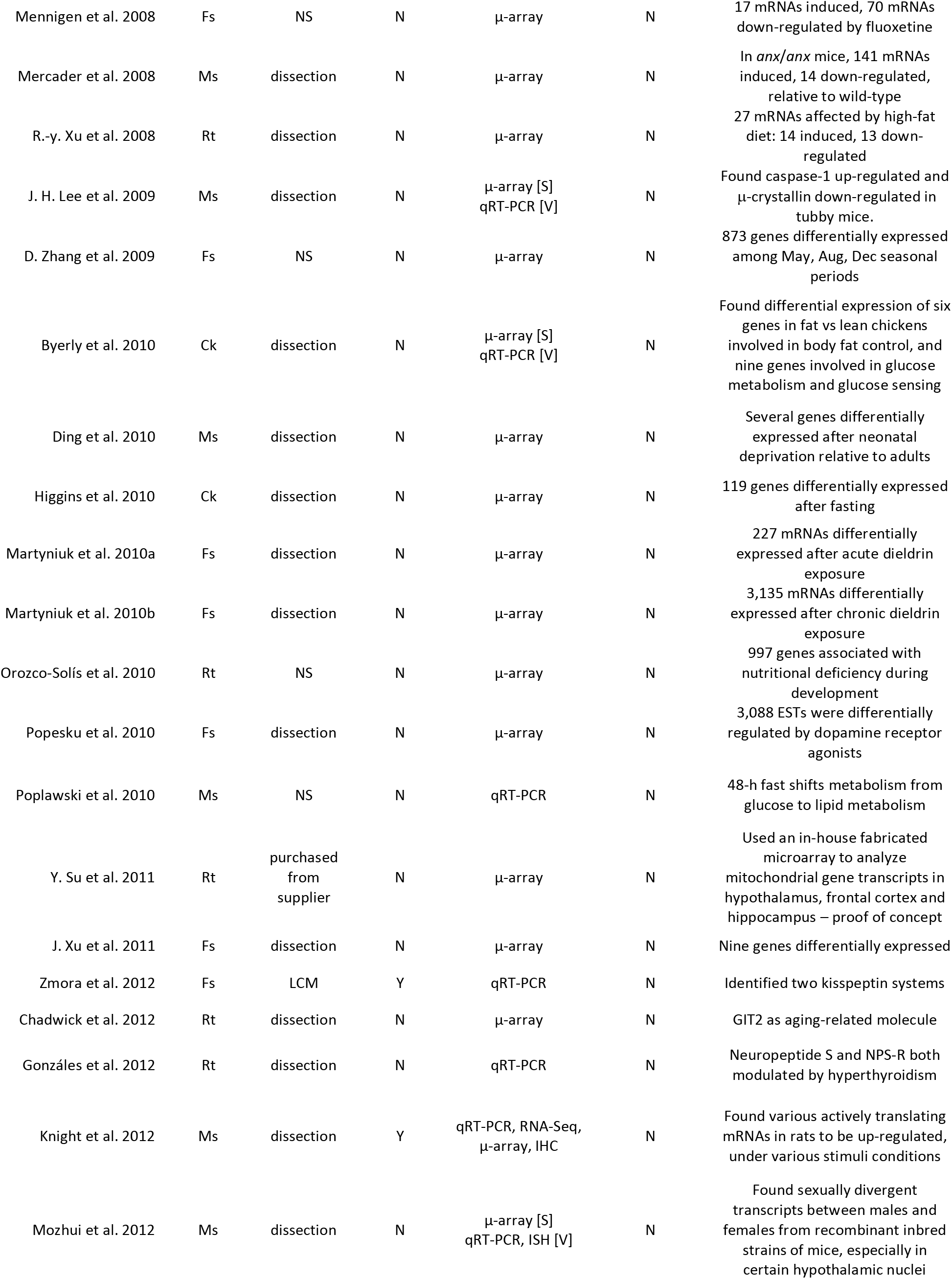
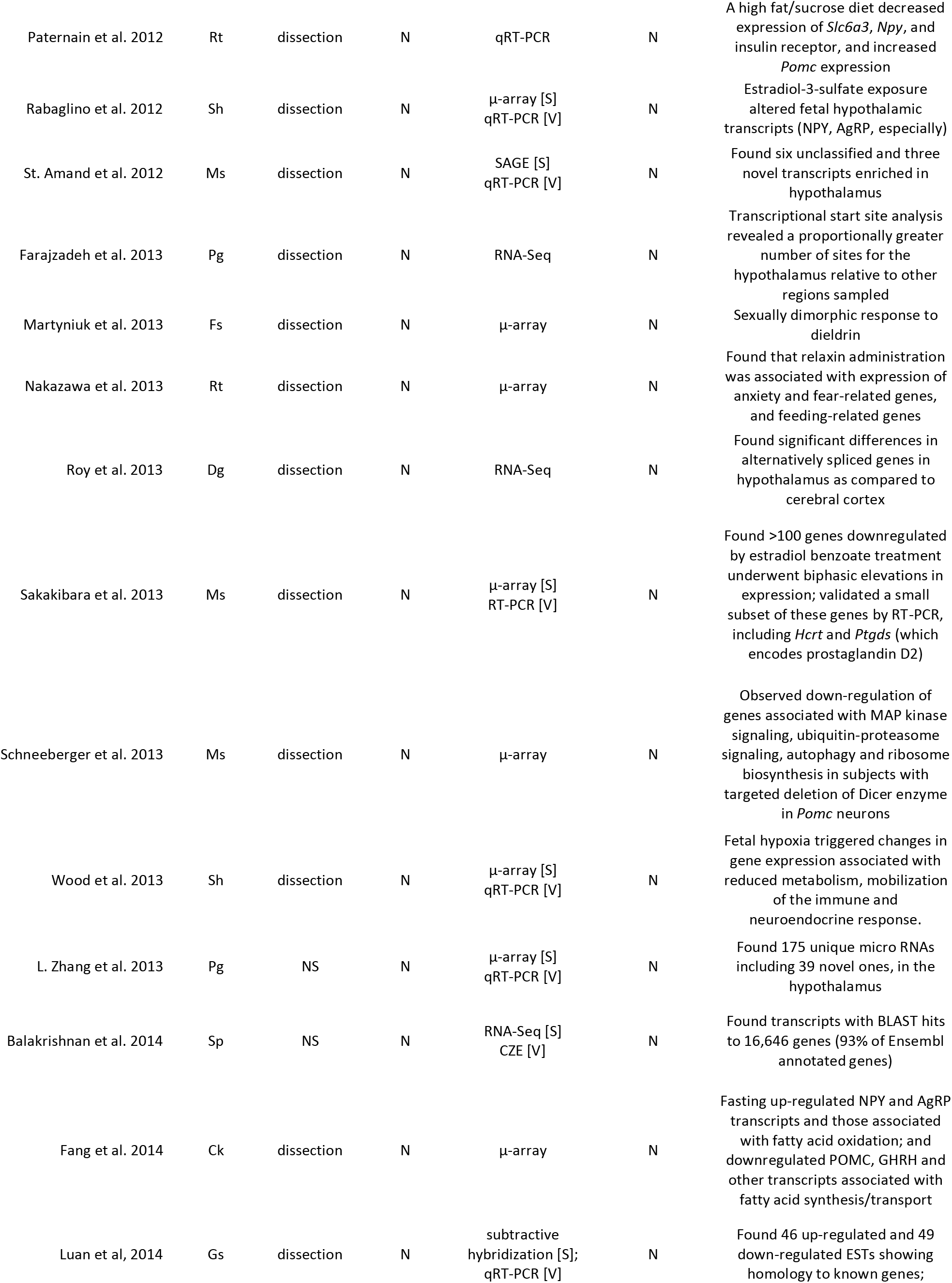
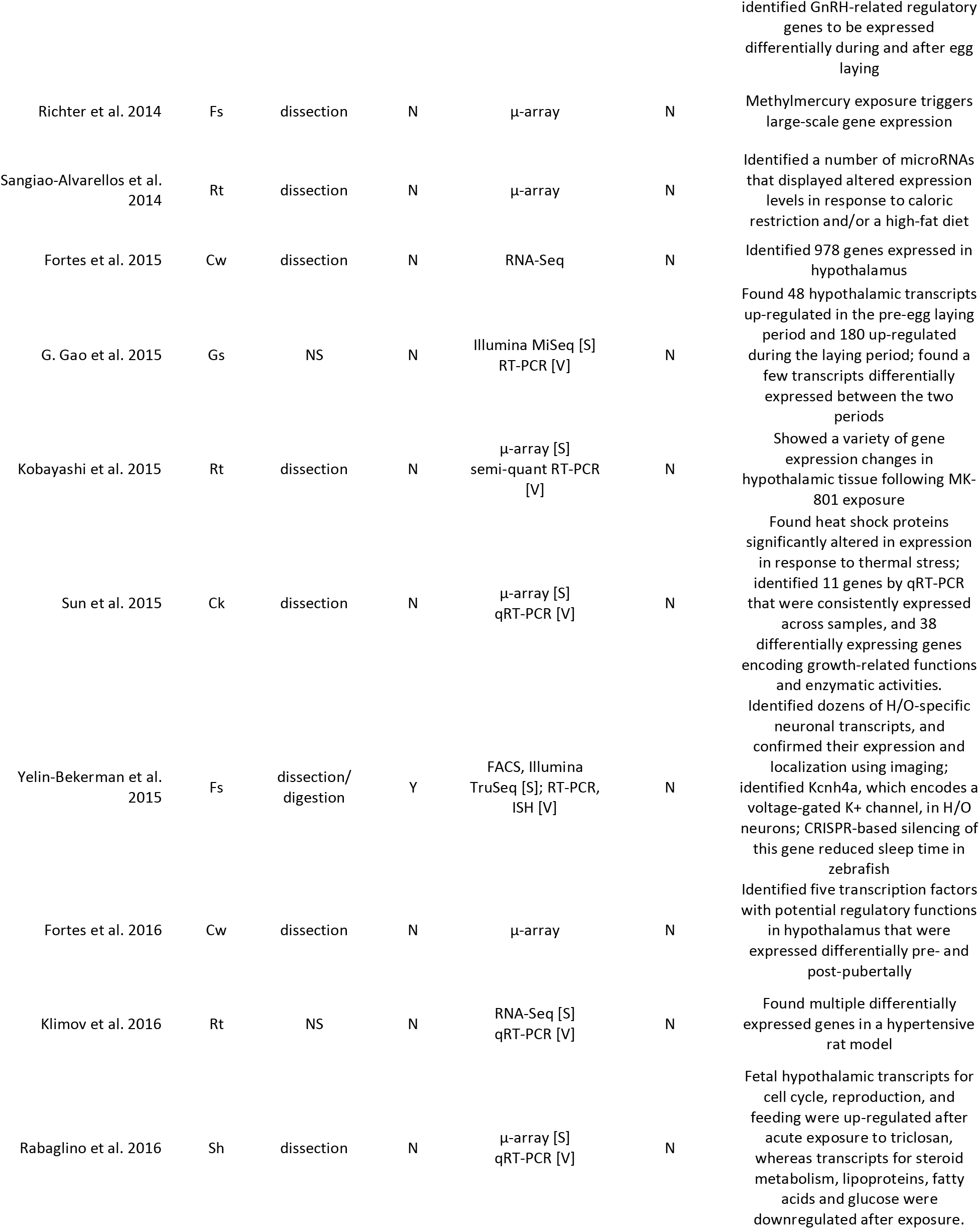
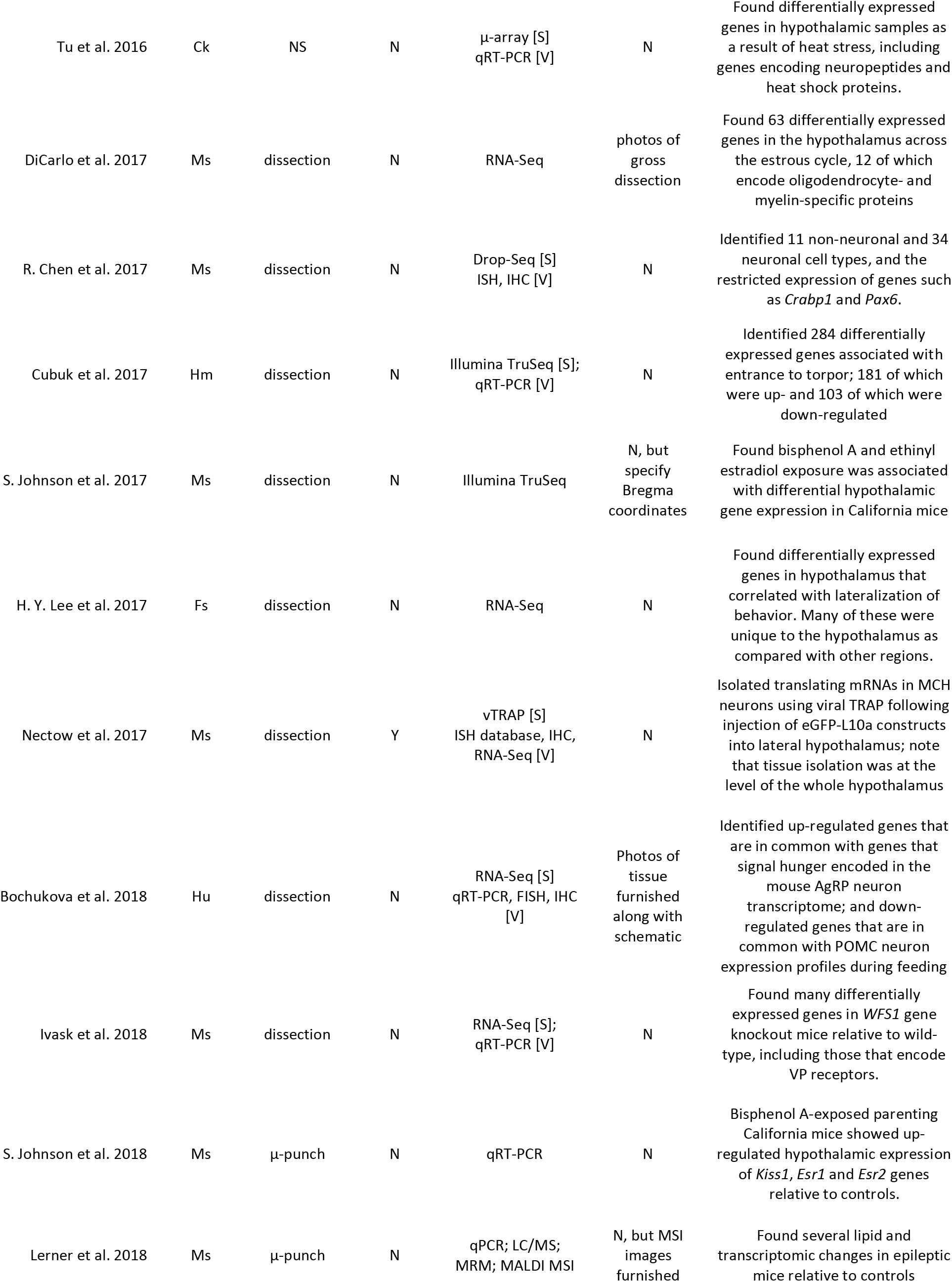
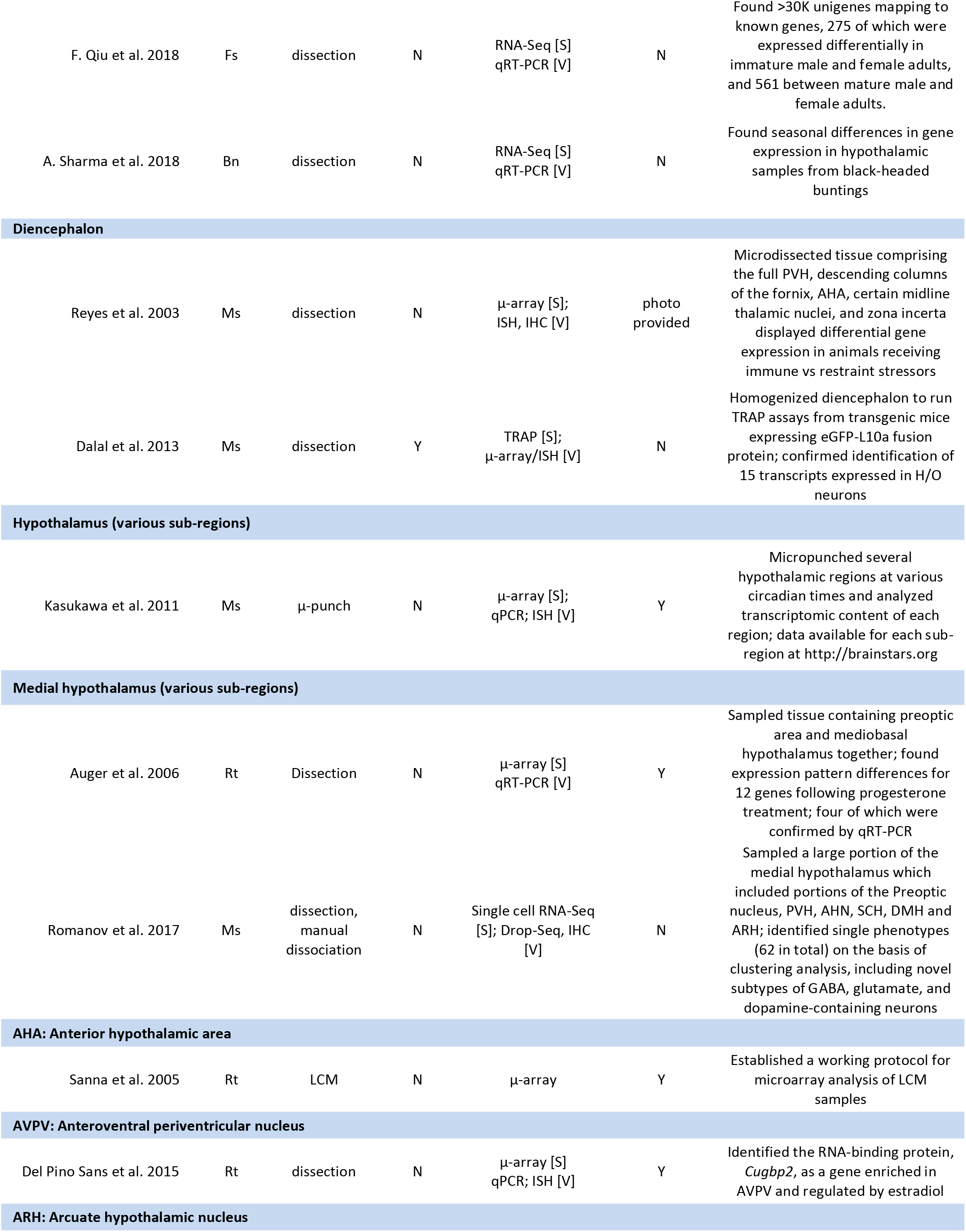
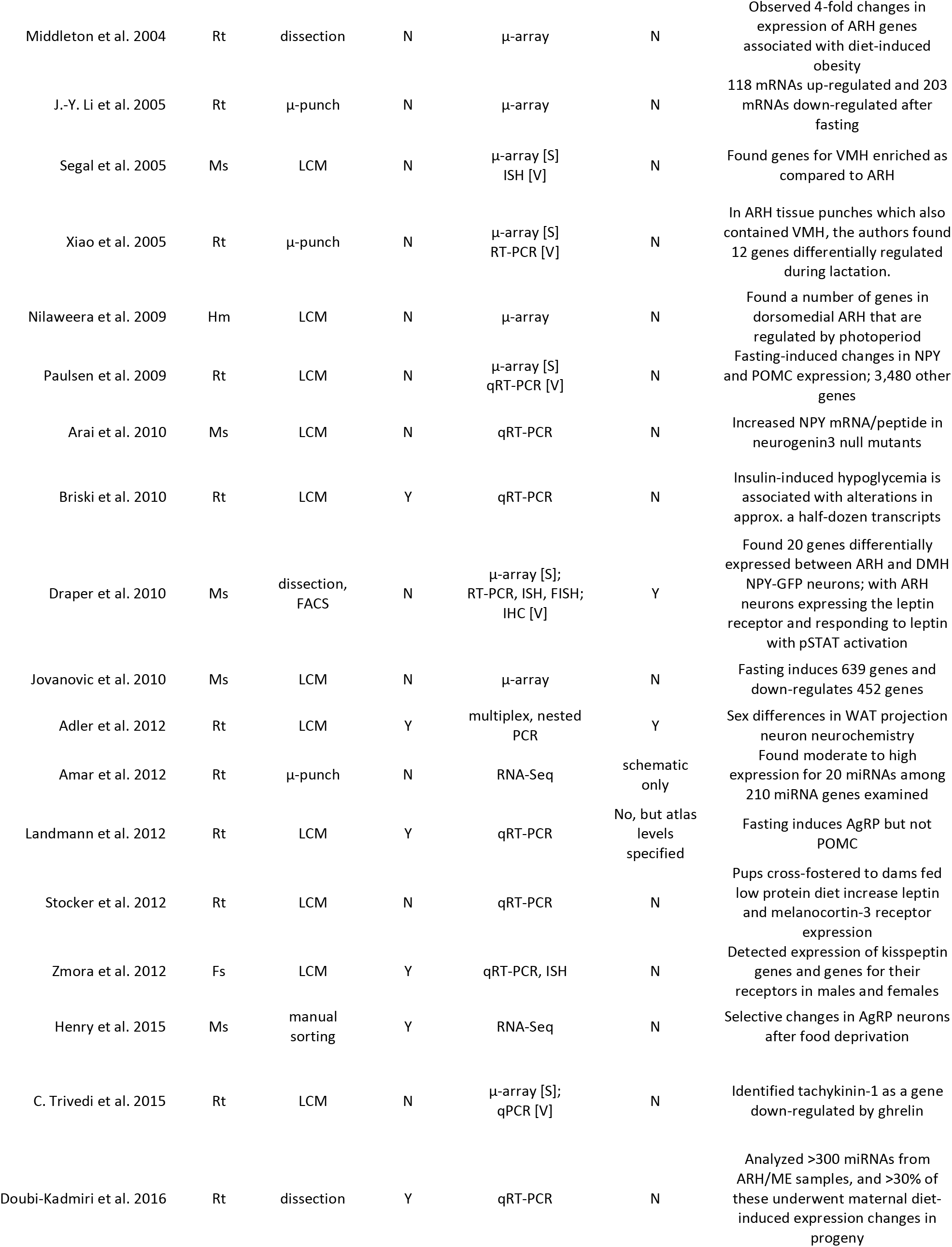
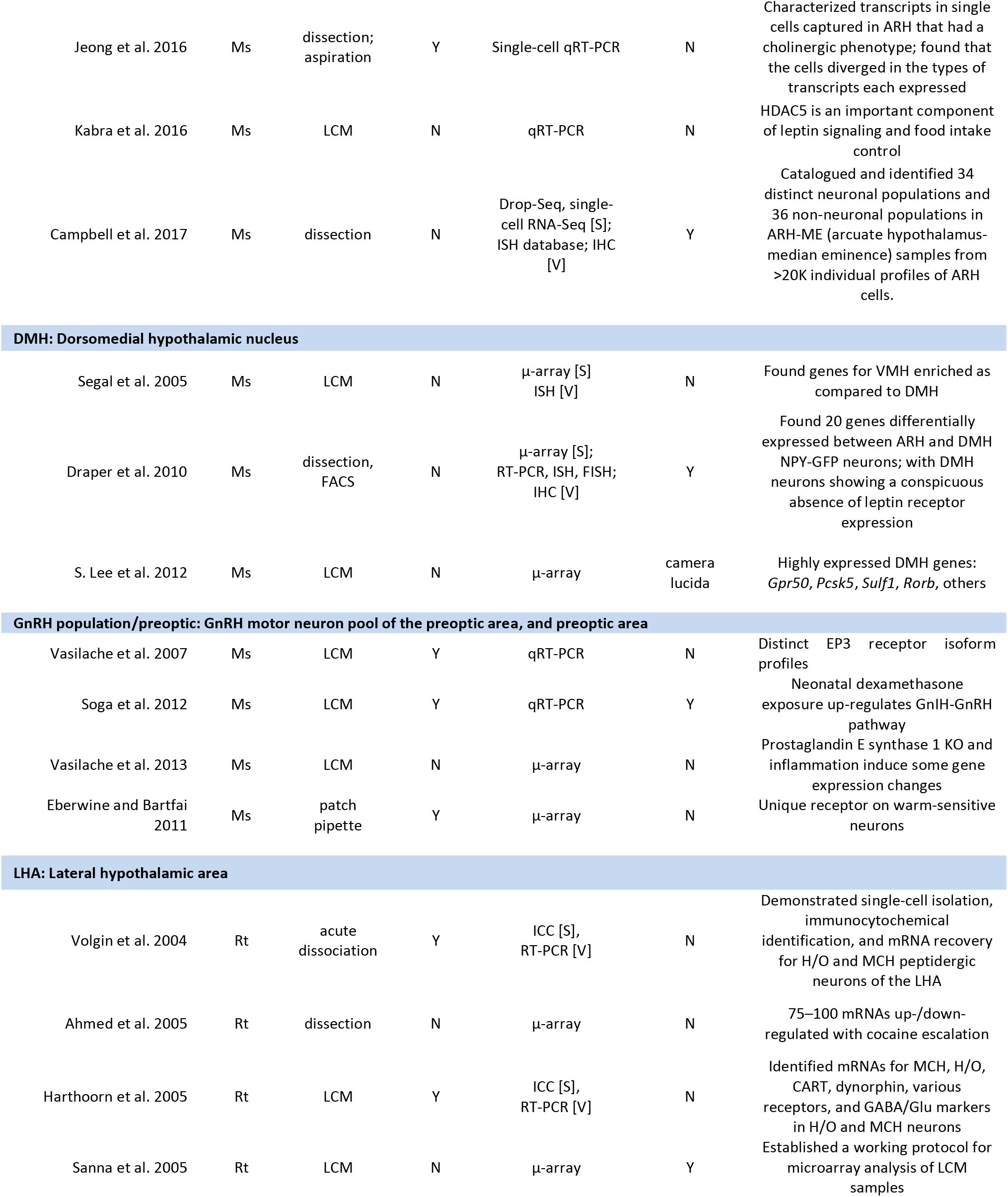
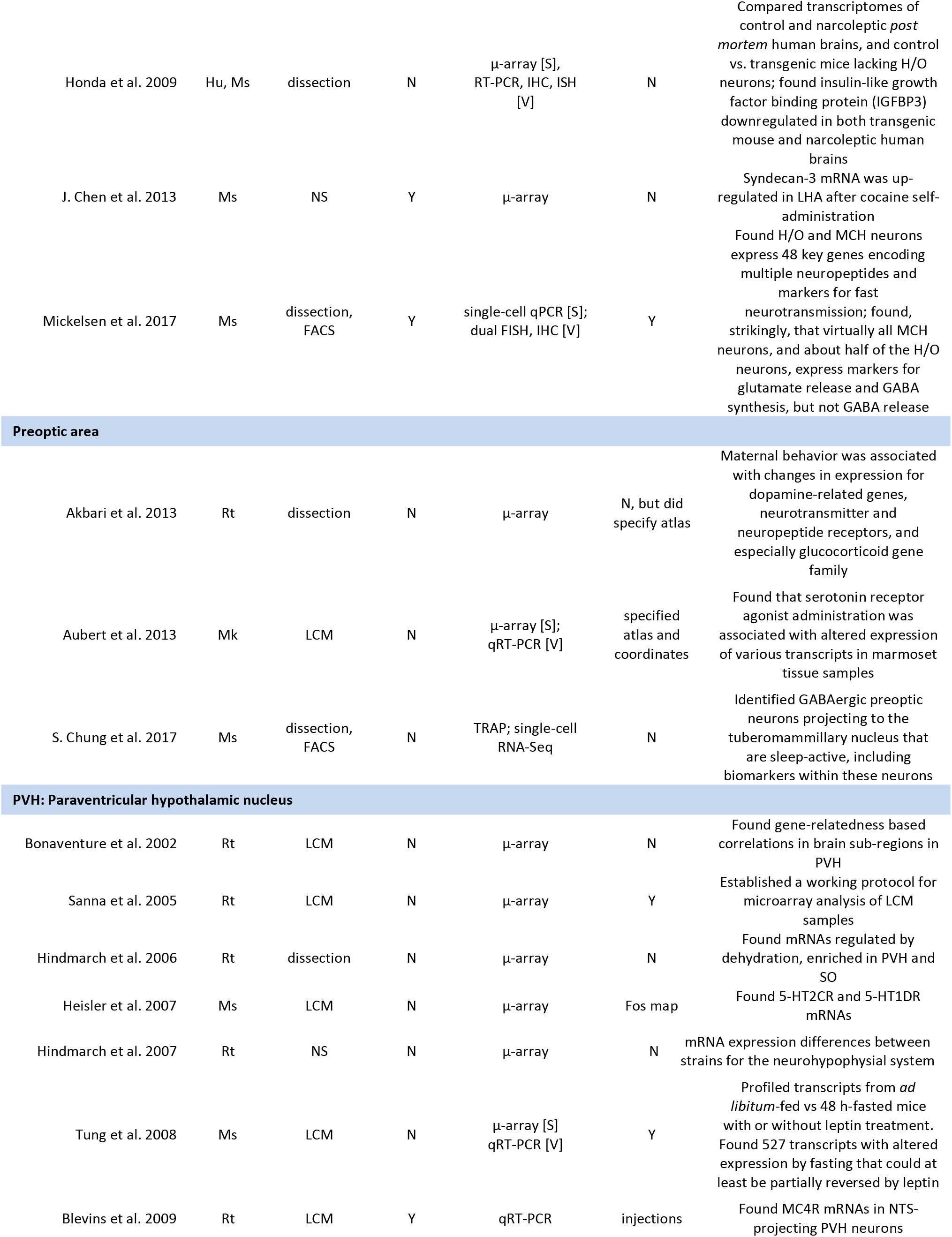
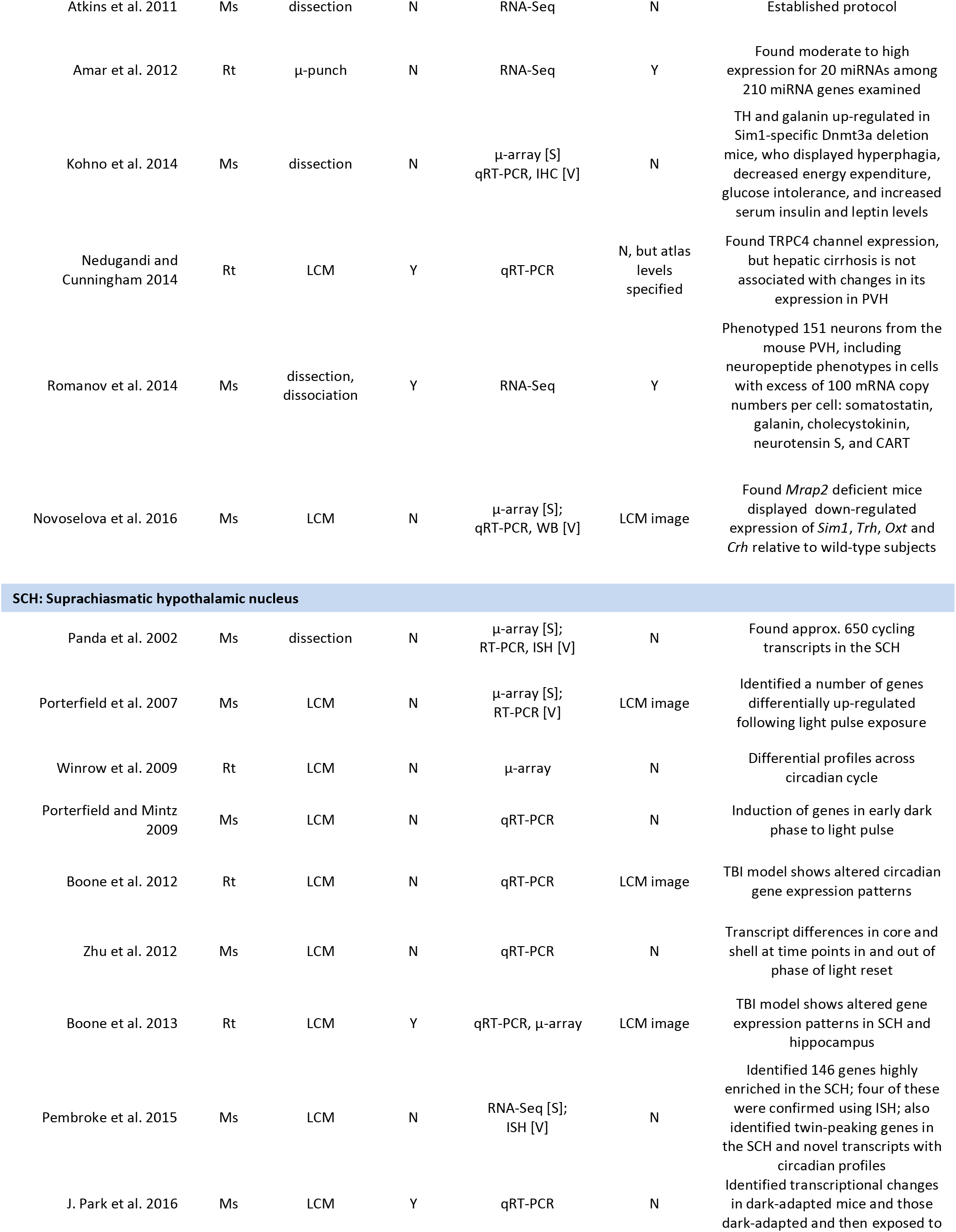
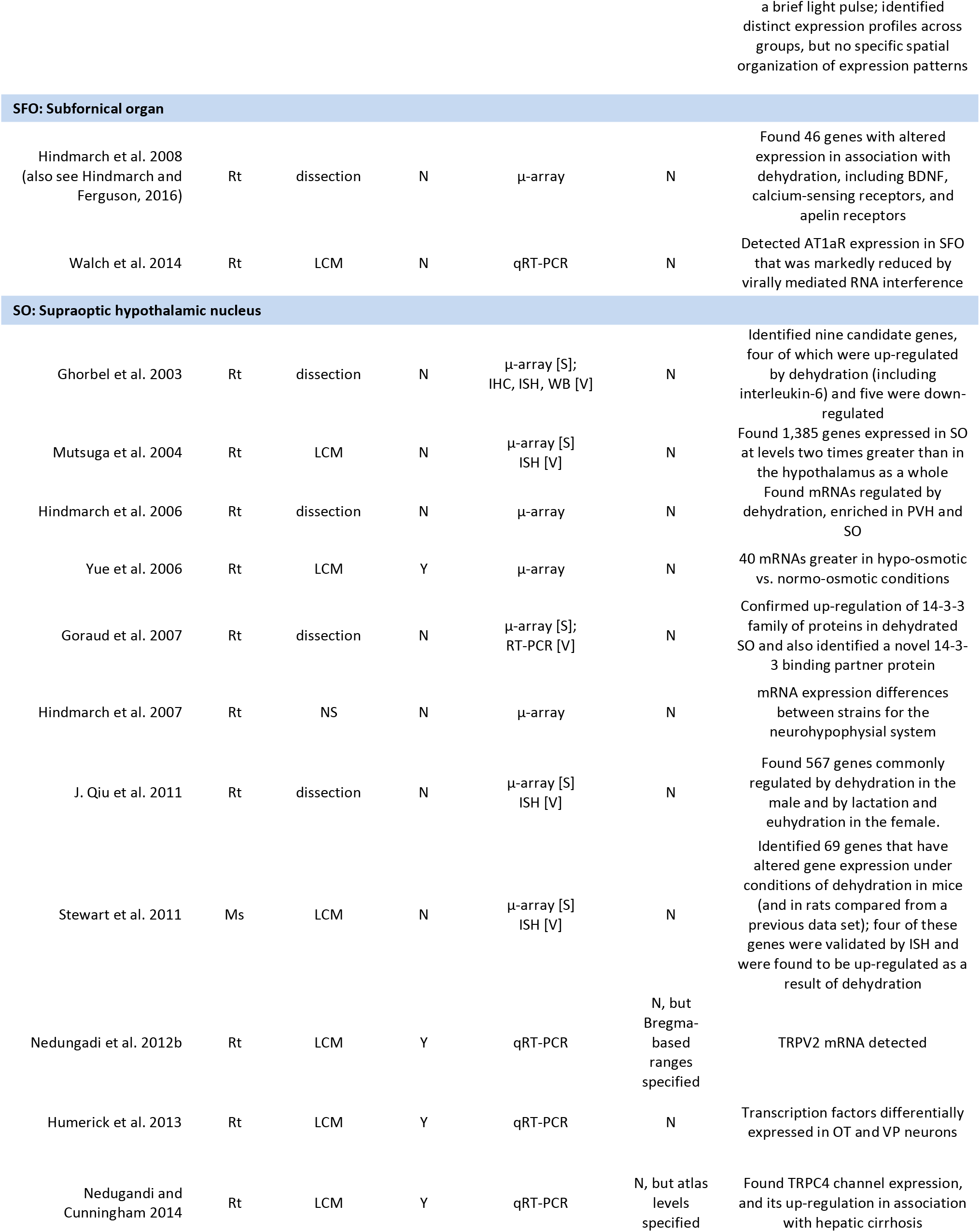
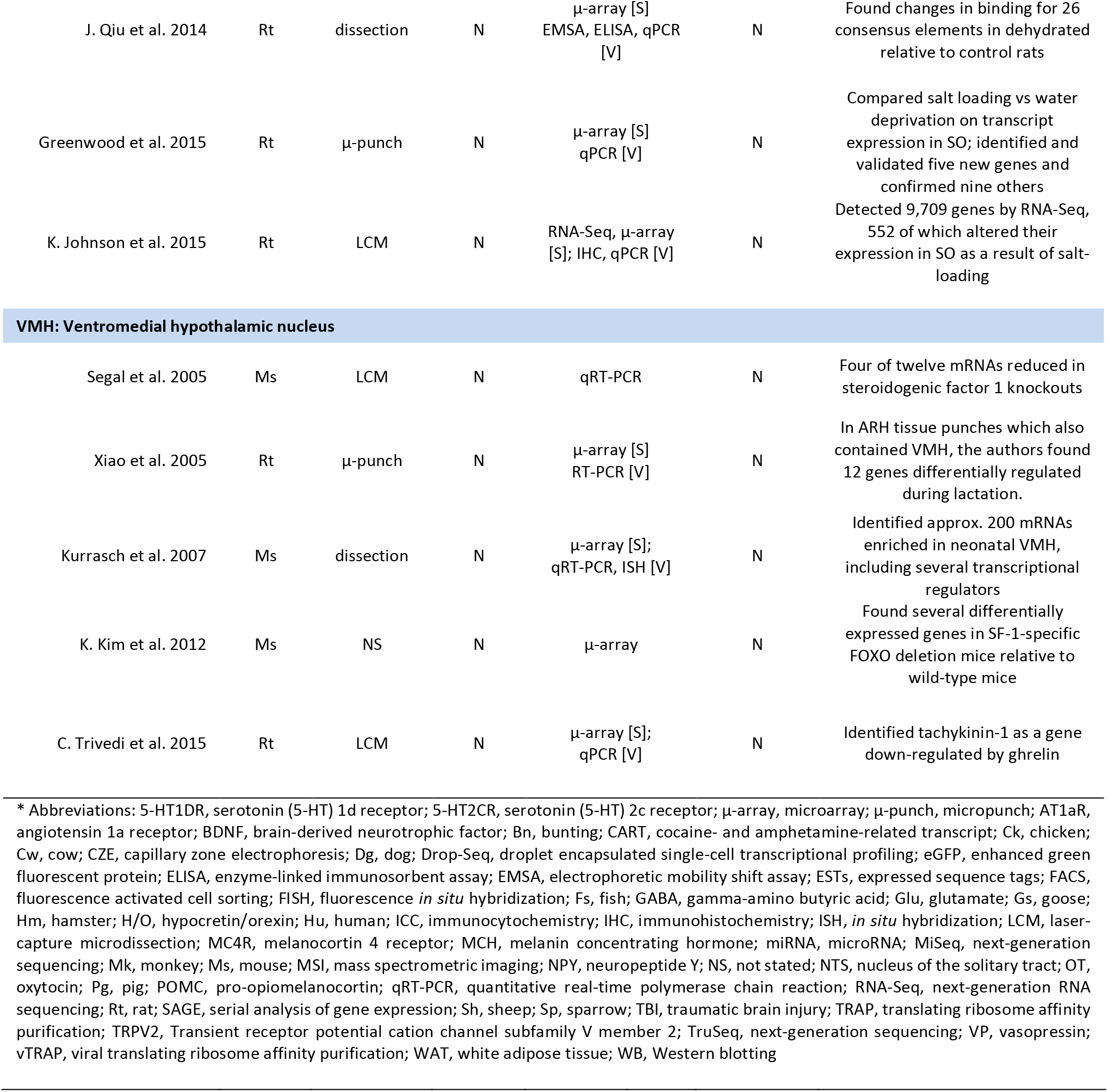
Selected transcriptomic studies in whole hypothalamus and by hypothalamic sub-region

**Table 2.**
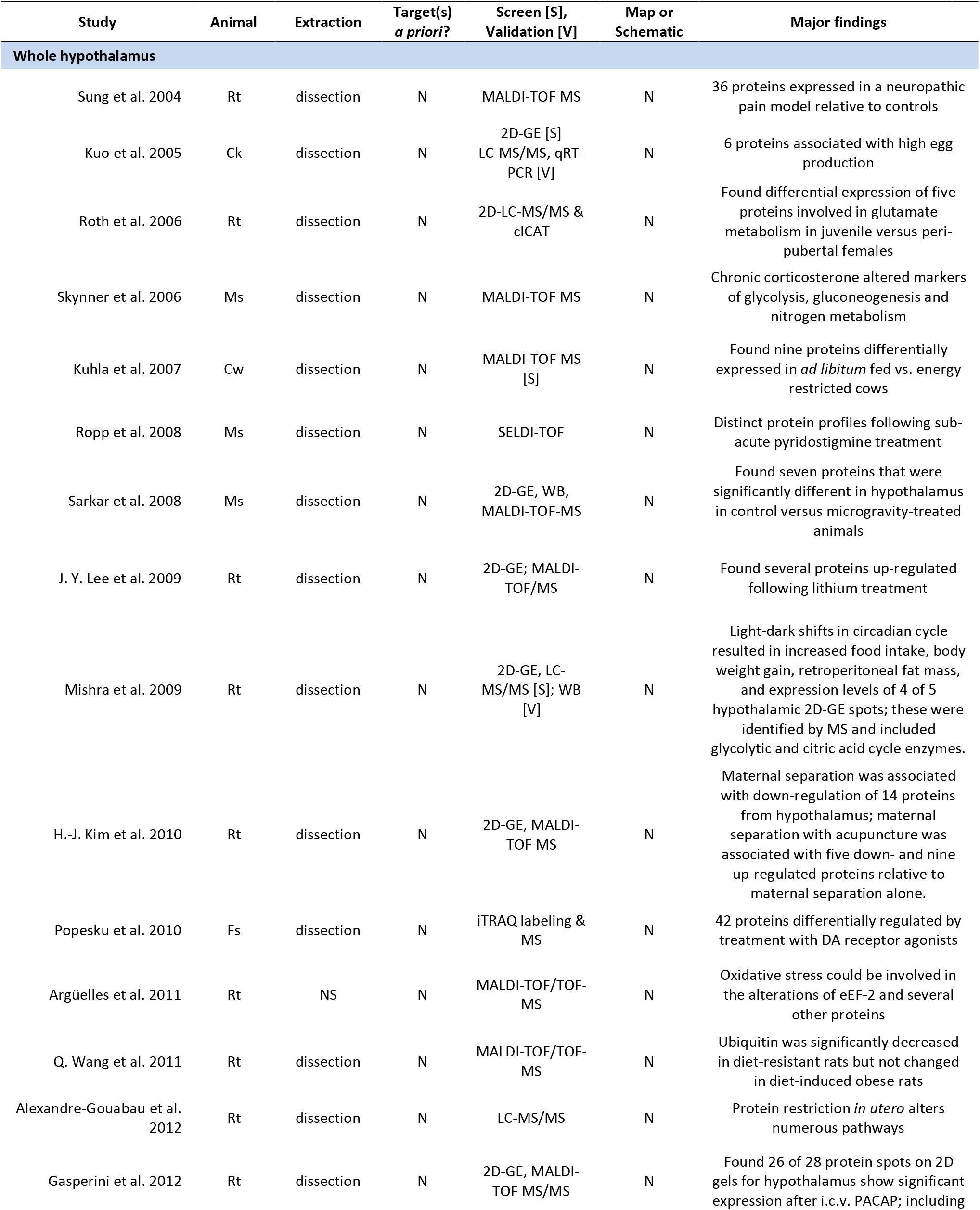
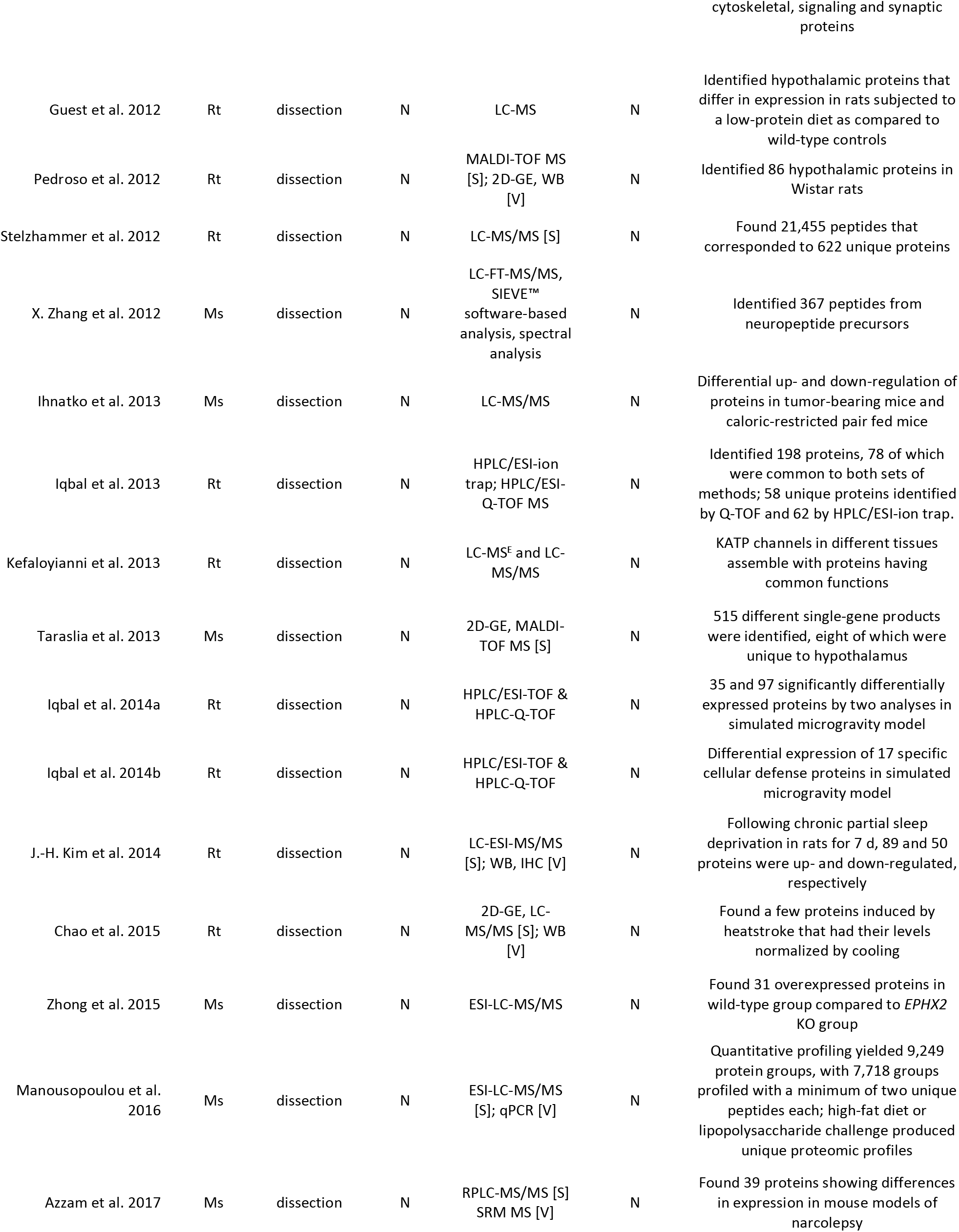
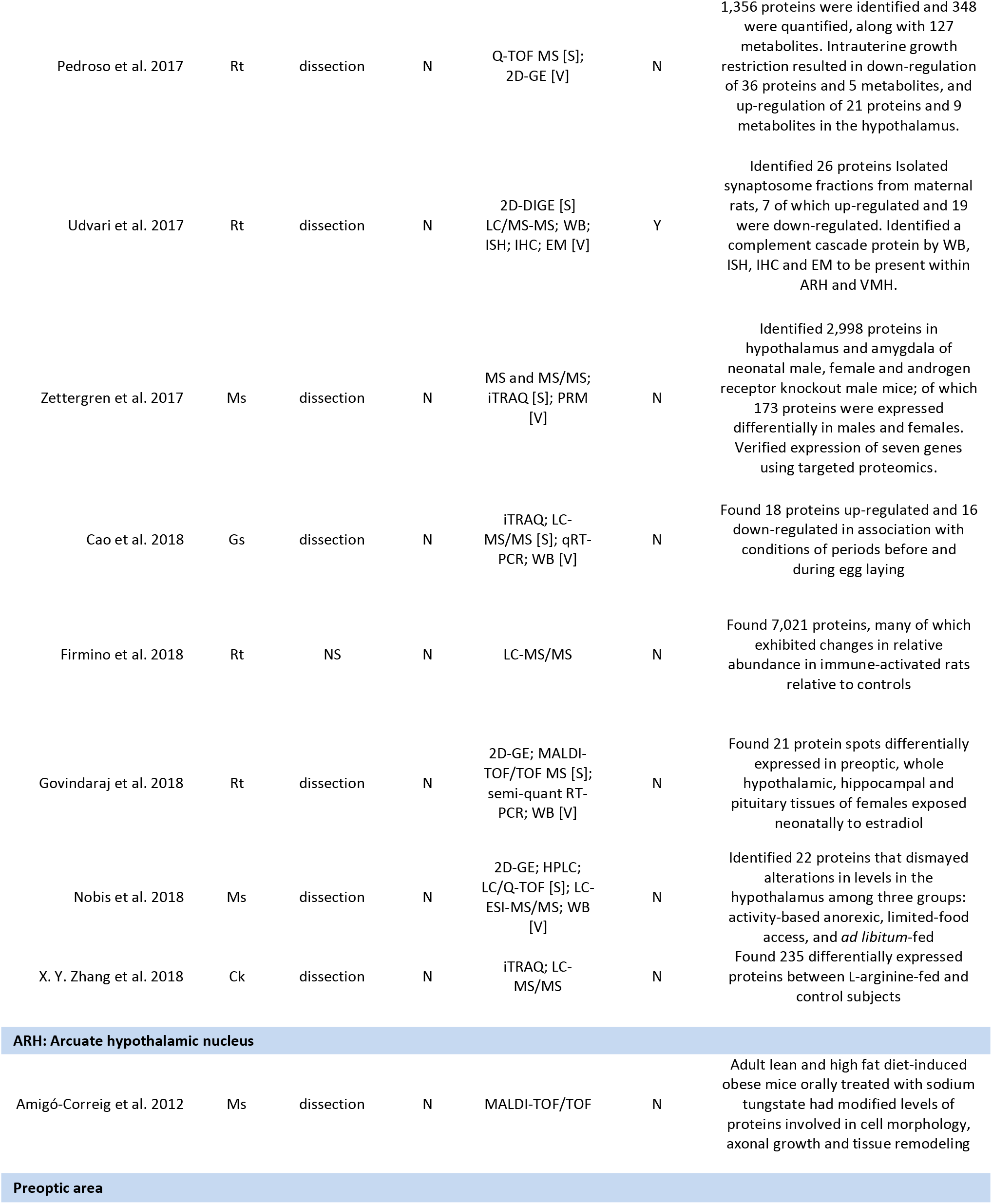
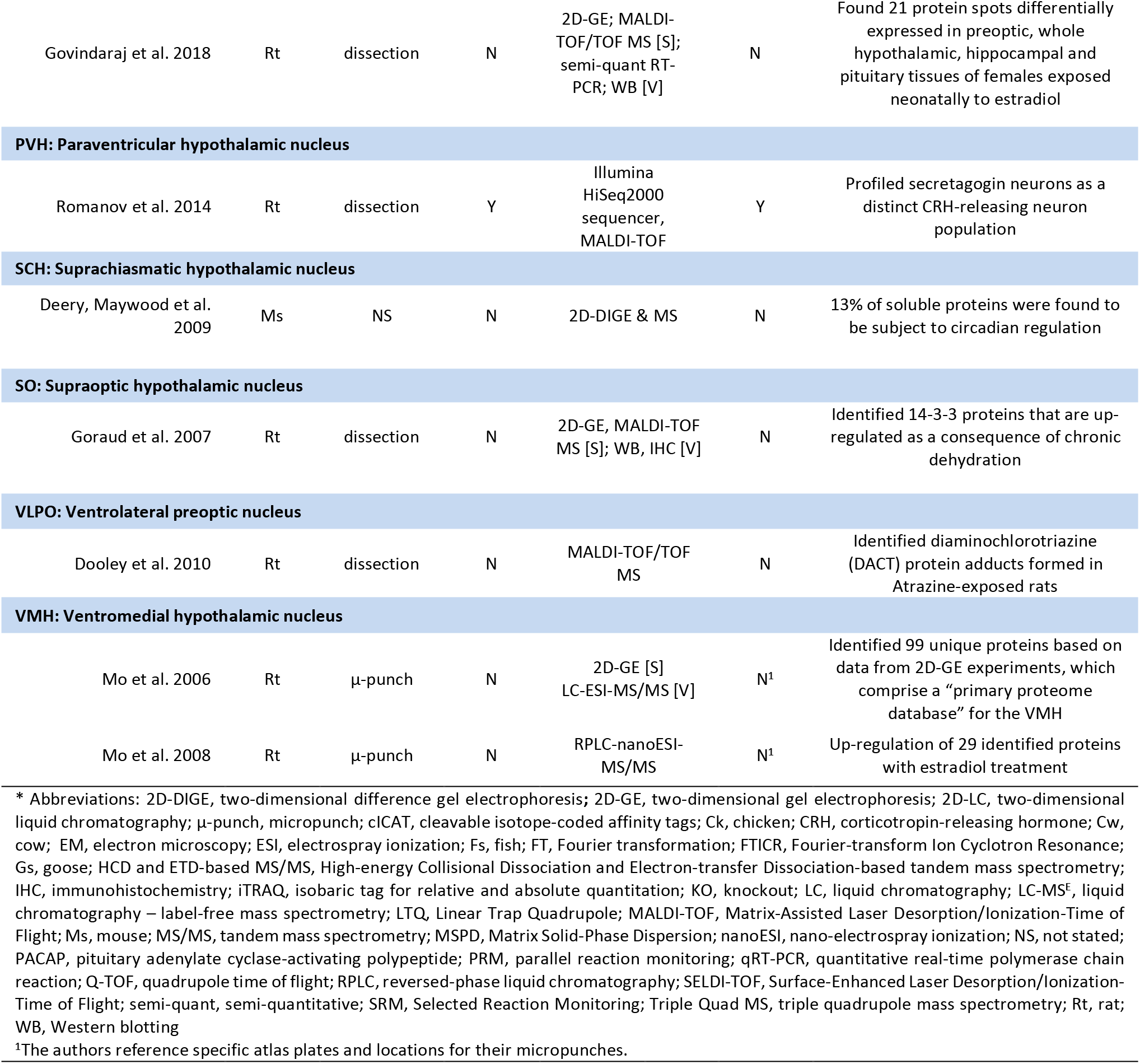
Selected proteomic studies in whole hypothalamus and by hypothalamic sub-region*

**Table 3.**
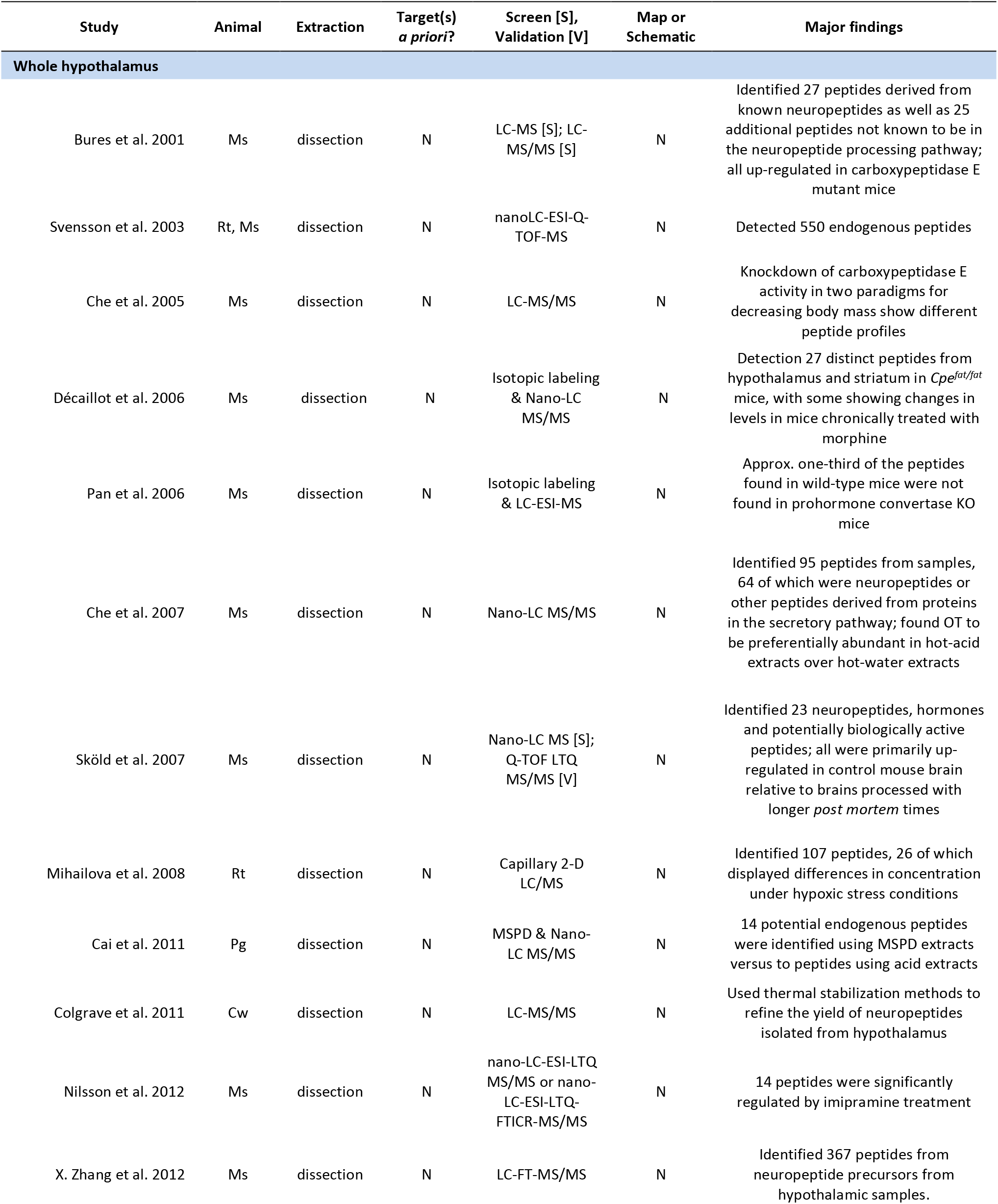
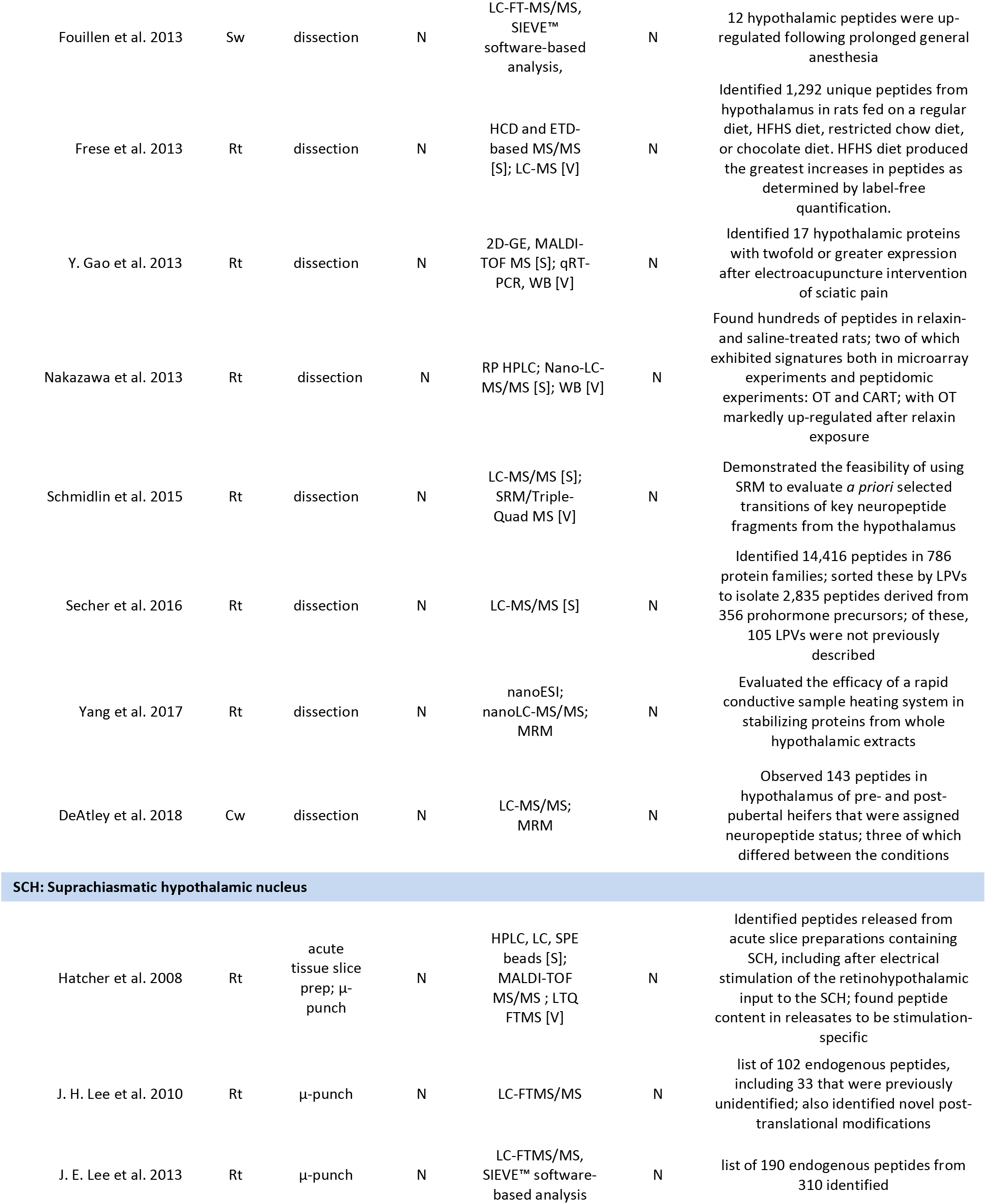
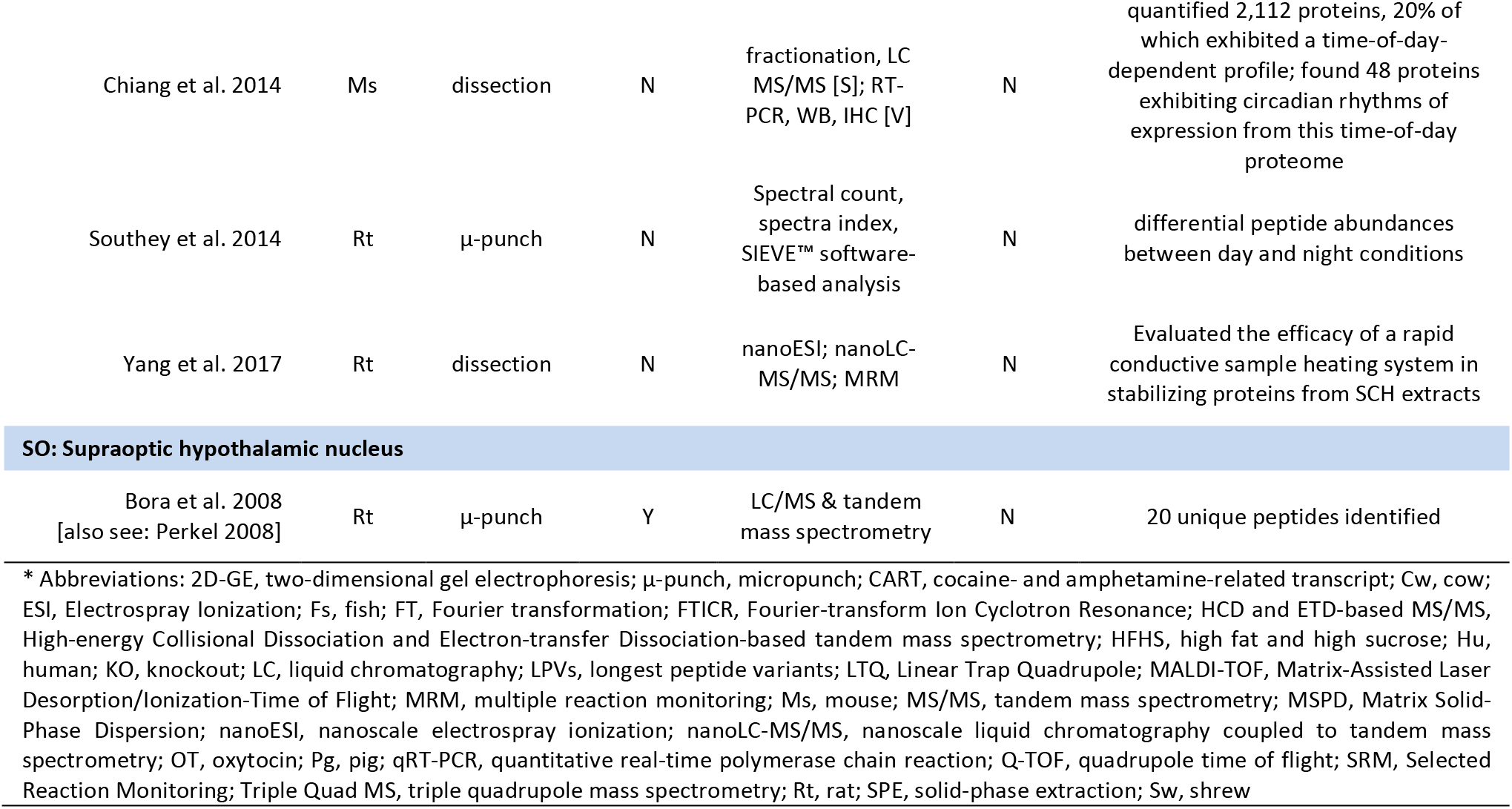
Selected peptidomic studies in whole hypothalamus and by hypothalamic sub-region*

| Study | Animal | Extraction | Target(s)<br>*a priori*? | Screen [S],<br>Validation [V] | Map or<br>Schematic | Major findings |
| --- | --- | --- | --- | --- | --- | --- |
| **Whole hypothalamus** |  |  |  |  |  |  |
| Bures et al. 2001 | Ms | dissection | N | LC-MS [S]; LC-MS/MS [S] | N | Identified 27 peptides derived from known neuropeptides as well as 25 additional peptides not known to be in the neuropeptide processing pathway; all up-regulated in carboxypeptidase E mutant mice |
| Svensson et al. 2003 | Rt, Ms | dissection | N | nanoLC-ESI-Q-TOF-MS | N | Detected 550 endogenous peptides |
| Che et al. 2005 | Ms | dissection | N | LC-MS/MS | N | Knockdown of carboxypeptidase E activity in two paradigms for decreasing body mass show different peptide profiles |
| Décaillot et al. 2006 | Ms | dissection | N | Isotopic labeling & Nano-LC MS/MS | N | Detection 27 distinct peptides from hypothalamus and striatum in *Cpe<sup>fat/fat</sup>* mice, with some showing changes in levels in mice chronically treated with morphine |
| Pan et al. 2006 | Ms | dissection | N | Isotopic labeling & LC-ESI-MS | N | Approx. one-third of the peptides found in wild-type mice were not found in prohormone convertase KO mice |
| Che et al. 2007 | Ms | dissection | N | Nano-LC MS/MS | N | Identified 95 peptides from samples, 64 of which were neuropeptides or other peptides derived from proteins in the secretory pathway; found OT to be preferentially abundant in hot-acid extracts over hot-water extracts |
| Sköld et al. 2007 | Ms | dissection | N | Nano-LC MS [S]; Q-TOF LTQ MS/MS [V] | N | Identified 23 neuropeptides, hormones and potentially biologically active peptides; all were primarily up-regulated in control mouse brain relative to brains processed with longer *post mortem* times |
| Mihailova et al. 2008 | Rt | dissection | N | Capillary 2-D LC/MS | N | Identified 107 peptides, 26 of which displayed differences in concentration under hypoxic stress conditions |
| Cai et al. 2011 | Pg | dissection | N | MSPD & Nano-LC MS/MS | N | 14 potential endogenous peptides were identified using MSPD extracts versus to peptides using acid extracts |
| Colgrave et al. 2011 | Cw | dissection | N | LC-MS/MS | N | Used thermal stabilization methods to refine the yield of neuropeptides isolated from hypothalamus |
| Nilsson et al. 2012 | Ms | dissection | N | nano-LC-ESI-LTQ MS/MS or nano-LC-ESI-LTQ-FTICR-MS/MS | N | 14 peptides were significantly regulated by imipramine treatment |
| X. Zhang et al. 2012 | Ms | dissection | N | LC-FT-MS/MS | N | Identified 367 peptides from neuropeptide precursors from hypothalamic samples. |
| Fouillen et al. 2013 | Sw | dissection | N | LC-FT-MS/MS, SIEVE™ software-based analysis, | N | 12 hypothalamic peptides were up-regulated following prolonged general anesthesia |
| Frese et al. 2013 | Rt | dissection | N | HCD and ETD-based MS/MS [S]; LC-MS [V] | N | Identified 1,292 unique peptides from hypothalamus in rats fed on a regular diet, HFHS diet, restricted chow diet, or chocolate diet. HFHS diet produced the greatest increases in peptides as determined by label-free quantification. |
| Y. Gao et al. 2013 | Rt | dissection | N | 2D-GE, MALDI-TOF MS [S]; qRT-PCR, WB [V] | N | Identified 17 hypothalamic proteins with twofold or greater expression after electroacupuncture intervention of sciatic pain |
| Nakazawa et al. 2013 | Rt | dissection | N | RP HPLC; Nano-LC-MS/MS [S]; WB [V] | N | Found hundreds of peptides in relaxin- and saline-treated rats; two of which exhibited signatures both in microarray experiments and peptidomic experiments: OT and CART; with OT markedly up-regulated after relaxin exposure |
| Schmidlin et al. 2015 | Rt | dissection | N | LC-MS/MS [S]; SRM/Triple-Quad MS [V] | N | Demonstrated the feasibility of using SRM to evaluate *a priori* selected transitions of key neuropeptide fragments from the hypothalamus |
| Secher et al. 2016 | Rt | dissection | N | LC-MS/MS [S] | N | Identified 14,416 peptides in 786 protein families; sorted these by LPVs to isolate 2,835 peptides derived from 356 prohormone precursors; of these, 105 LPVs were not previously described |
| Yang et al. 2017 | Rt | dissection | N | nanoESI; nanoLC-MS/MS; MRM | N | Evaluated the efficacy of a rapid conductive sample heating system in stabilizing proteins from whole hypothalamic extracts |
| DeAtley et al. 2018 | Cw | dissection | N | LC-MS/MS; MRM | N | Observed 143 peptides in hypothalamus of pre- and post-pubertal heifers that were assigned neuropeptide status; three of which differed between the conditions |
| SCH: Suprachiasmatic hypothalamic nucleus |  |  |  |  |  |  |
| Hatcher et al. 2008 | Rt | acute tissue slice prep; μ-punch | N | HPLC, LC, SPE beads [S]; MALDI-TOF MS/MS ; LTQ FTMS [V] | N | Identified peptides released from acute slice preparations containing SCH, including after electrical stimulation of the retinohypothalamic input to the SCH; found peptide content in releasates to be stimulation-specific |
| J. H. Lee et al. 2010 | Rt | μ-punch | N | LC-FTMS/MS | N | list of 102 endogenous peptides, including 33 that were previously unidentified; also identified novel post-translational modifications |
| J. E. Lee et al. 2013 | Rt | μ-punch | N | LC-FTMS/MS, SIEVE™ software-based analysis | N | list of 190 endogenous peptides from 310 identified |
| Chiang et al. 2014 | Ms | dissection | N | fractionation, LC MS/MS [S]; RT-PCR, WB, IHC [V] | N | quantified 2,112 proteins, 20% of which exhibited a time-of-day-dependent profile; found 48 proteins exhibiting circadian rhythms of expression from this time-of-day proteome |
| Southey et al. 2014 | Rt | μ-punch | N | Spectral count, spectra index, SIEVE™ software-based analysis | N | differential peptide abundances between day and night conditions |
| Yang et al. 2017 | Rt | dissection | N | nanoESI; nanoLC-MS/MS; MRM | N | Evaluated the efficacy of a rapid conductive sample heating system in stabilizing proteins from SCH extracts |

**SO: Supraoptic hypothalamic nucleus**

|  |  |  |  |  |  |  |
| --- | --- | --- | --- | --- | --- | --- |
| Bora et al. 2008<br>[also see: Perkel 2008] | Rt | μ-punch | Y | LC/MS & tandem mass spectrometry | N | 20 unique peptides identified |

\* Abbreviations: 2D-GE, two-dimensional gel electrophoresis; μ-punch, micropunch; CART, cocaine- and amphetamine-related transcript; Cw, cow; ESI, Electrospray Ionization; Fs, fish; FT, Fourier transformation; FTICR, Fourier-transform Ion Cyclotron Resonance; HCD and ETD-based MS/MS, High-energy Collisional Dissociation and Electron-transfer Dissociation-based tandem mass spectrometry; HFHS, high fat and high sucrose; Hu, human; KO, knockout; LC, liquid chromatography; LPVs, longest peptide variants; LTQ, Linear Trap Quadrupole; MALDI-TOF, Matrix-Assisted Laser Desorption/Ionization-Time of Flight; MRM, multiple reaction monitoring; Ms, mouse; MS/MS, tandem mass spectrometry; MSPD, Matrix Solid-Phase Dispersion; nanoESI, nanoscale electrospray ionization; nanoLC-MS/MS, nanoscale liquid chromatography coupled to tandem mass spectrometry; OT, oxytocin; Pg, pig; qRT-PCR, quantitative real-time polymerase chain reaction; Q-TOF, quadrupole time of flight; SRM, Selected Reaction Monitoring; Triple Quad MS, triple quadrupole mass spectrometry; Rt, rat; SPE, solid-phase extraction; Sw, shrew

A number of observations can be made from an examination of the figure. First, of the total number of studies listed in **Tables 1–3**, 45–83% of them (depending on which molecular analysis was performed) provide no sub-regional specificity for their sampling but rather sample the whole hypothalamus (**Fig. 3A**). Second, of the studies performing high throughput extraction and molecular analysis of hypothalamic sub-regions, the greatest degree of coverage occurs for transcriptomic (**Fig. 3B**), followed by proteomic (**Fig. 3C**) and peptidomic (**Fig. 3D**) studies. Finally, across all methods, the overwhelming emphasis of sub-regional analyses of the hypothalamus has been on medially located nuclei, with little to no examination of sub-regions within the larger lateral hypothalamic area (LHA). Even for transcriptomic studies (**Fig. 3B**), the greater majority of studies of the LHA have focused mainly on a few key peptidergic cell types and not the whole region *per se*. Below, after describing a few studies that have focused on the hypothalamus in the context of whole-brain or multi-regional studies, we summarize a few key studies from among those listed in **Tables 1–3**.

#### 4.2.1: Whole brain extraction and multi-region comparison studies

There are many excellent reasons investigators opt to extract molecular information from the whole brain or large subdivisions of the brain without attending to where exactly in the brain the molecules are located. Such reasons include the need for investigators to survey the effects of factors that produce global, whole-organism or whole-subdivision effects that are poorly understood at a regional or cellular level. These include environmental agents (X. Li et al. 2014), pharmacological interventions (Jin et al. 2016), ontogenic state (e.g., see introductory remarks in Balivada et al. 2017), or physiological processes. J. Miller et al. (2014) examined various hypothalamic sub-regions within the context of a hemispheric tissue analysis in prenatal human brain using high throughput transcriptomic methods. Zapala et al. (2005) contextualized regional specificity with embryonic development, taking care to provide supplementary information that includes photographic documentation of the tissue they dissected for their hypothalamic sample. In contrast, it is disappointing that in their “in-depth analysis of the mouse brain and its major regions and cell types” for the proteome, K. Sharma et al. (2015) neglected to sample the hypothalamus in what is otherwise a detailed and interesting study.

#### 4.2.2: Molecular extraction from whole hypothalamus

In non-mammalian vertebrates, the hypothalamus has been studied for transcriptomics, proteomics, and peptidomics in fishes and birds; in some cases, in the context of animal husbandry. For example, hypothalamic and pituitary molecules associated with high egg production in chickens have been analyzed at the transcriptomic (Shiue et al. 2006; L.-R. Chen et al. 2007; **Table 1**) and proteomic (Kuo et al. 2005) levels. Egg-laying traits have also been compared alongside transcripts identified to be associated with high egg production (C.-F. Chen et al. 2007). The hypothalamic transcriptome and proteome of the Huoyan goose (Luan et al. 2014; Cao et al. 2018) and the hypothalamic transcriptome of Sichuan white goose (G. Gao et al. 2015) have been profiled before, during, or after their egg laying periods in the interests of finding clues to improve the reproductive performance of these economically valuable domestic animals (also see Figure 1 of H. F. Li et al. 2011). In the interests of optimizing feed intake in chickens or to understand how they cope with environmentally-induced pressures, many studies have also examined the role of body composition, fasting, diet, or heat stress on gene expression in chicken hypothalamus (e.g., Byerly et al. 2010; Fang et al. 2014; Sun et al. 2015; Tu et al. 2016; see also Kuenzel et al. 1999). Despite the intensive investigations of chicken hypothalamus for molecular mining and extraction, these studies have not contextualized sub-regional changes in expression for molecules in relation to published stereotaxic atlases of the chicken that include illustrations, maps and drawings of the hypothalamus with stereotaxic coordinates (van Tienhoven and Juhász 1962; Yasuda and Lepkovsky 1969; Feldman et al. 1973). Seasonal changes in hypothalamic gene expression have also been documented in the black-headed bunting, a migratory songbird (A. Trivedi et al. 2014; A. Sharma et al. 2018).

In mammals, whole hypothalamus has been mined for gene transcripts in mouse, rat, hamster, guinea pig, shrew pig, cow, sheep, dog and human (**Table 1**). Recently, human induced pluripotent stem cells differentiated into “hypothalamic-like” neurons have also been profiled for their transcriptomes (Rajamani et al. 2018). The first large-scale *in situ* hybridization-based study of hypothalamus-enriched transcripts was provided by Gautvik et al. (1996) in the rat by using directional tag PCR subtraction, which led to the discovery of the hypocretin neuropeptides (de Lecea et al., 1998; also see Sutcliffe, 2001; Sutcliffe and de Lecea, 2002). Friedman and colleagues utilized a novel molecular technique that extends the principles underlying an earlier approach (Heiman et al. 2008), to isolate and extract activated transcriptional systems in the hypothalamus under conditions of salt-loading, fasting, light exposure, or various other stimulus paradigms (Knight et al. 2012). Specifically, they immunoprecipitated the phosphorylated form of the ribosomal protein, S6, to isolate and enrich mRNAs that are actively being translated (i.e., in transcriptionally activated neurons) in mouse hypothalamic samples. Using TaqMan^®^ technology (Holland et al. 1991), RNA-Seq and microarrays, they isolated several mRNAs, many of which displayed expression in pS6-immunoreactive neurons in various sub-regions of the hypothalamus.

Using Drop-Seq, a method that allows for single-cell transcriptomics to be performed in a manner that preserves the cell provenance of the RNA that is extracted (Macosko et al. 2015), R. Chen et al. (2017) reported single-cell RNA sequencing results from the adult mouse hypothalamus. They used clustering analysis to identify 11 non-neuronal (including oligodendrocytes, astrocytes, ependymocytes, tanycytes, microglia, and macrophages) and 34 neuronal cell types (including 15 glutamatergic and 18 GABAergic clusters, and one histaminergic neuron cluster) from tissue dissociated from manually dissected hypothalamus, and confirmed some of their key findings by performing immunohistochemistry for neuropeptides or comparing their results with those found in the publicly available Allen Brain Atlas. Importantly, their workflow revealed the spatially restricted expression of novel molecules in the hypothalamus, including retinoic acid binding protein (*Crabp1*) in the ARH. They also found restricted expression of the neurodevelopmental factor, *Pax6*, in the zona incerta, which the authors assign as a hypothalamic structure but which is considered as a thalamic structure by others (e.g., see Swanson 2018). Importantly, their datasets indicate that all hypothalamic peptidergic neurons can also be classified by the small neurotransmitter they synthesize (glutamate or GABA). Recently, Romanov et al. (2017) provided evidence of numerous novel neuronal phenotypes of hypothalamic cells using single cell RNA-Seq and DropSeq technologies, but the only provenance that could be attributed to these cells was from within the large heterogeneous group of hypothalamic subregions partially sampled within their microdissected tissue sample, which include large portions of the medial, but not lateral hypothalamus. In contrast, Yelin-Bekerman et al. (2015) sampled from the whole hypothalamus of zebrafish to identify transcripts specific to neurons – isolated by fluorescence-activated cell sorting (FACS) – that expressed the neuropeptide hypocretin/orexin (H/O); these neurons are typically enriched in the lateral hypothalamus in most species (**Table 1**).

To date, few studies have examined proteomic or peptidomic profiles of whole hypothalamic samples. Extending the protocol they developed for peptidomic analysis of small microdissected brain regions such as the motor cortex, thalamus and striatum (Sköld et al. 2002), the Andrén laboratory reported identifying novel peptides from hypothalamic extracts (Svensson et al. 2003; Sköld et al. 2007). Fälth et al. (2006) developed a database for endogenous peptides identified by mass spectrometry, into which they have incorporated their hypothalamic datasets. Nakazawa et al. (2013) took the rather novel approach of performing both transcriptomics and peptidomics on separate sets of whole hypothalamic extracts (a “cross-omics” approach), and reported consensus results from both methods for oxytocin up-regulation in association with intracerebroventricular relaxin administration in rats. Recently, “cross-omics” approaches have been extended to combined transcriptomics/lipidomics of hypothalamus (Lerner et al. 2018).

#### 4.2.3: Molecular extraction from the hypothalamic circadian system

The suprachiasmatic hypothalamic nucleus (SCH), a well-defined compact nucleus within the hypothalamus that is amenable to precise sampling or molecular studies (e.g., see Fig. 1 of Porterfield et al. 2007; Fig. 1 of Boone et al. 2013), is the primary neural substrate for the master circadian clock in the body, which receives signals that allow organisms to respond to shifts in light during the day-night cycle. Often, circadian rhythms are characterized by changes in gene expression within the SCH; studies using microarray analysis demonstrated, for example, that approximately 650 transcripts undergo cyclic changes in expression in the SCH and the liver of mice, with many of these specific to the SCH (Panda et al. 2002). After certain stimuli, immediate early genes in the SCH peak and return to baseline, while a few others maintain their expression levels to protect the nuclei from excitotoxicity (Porterfield et al. 2007; Porterfield and Mintz 2009). Similar to contrasts between light and dark cycles, the transcriptome of the SCH is also distinct during wake and sleep cycles (Winrow et al. 2009), and there is recent transcriptomic evidence that certain classes of genes in the SCH peak twice in their expression levels across the circadian cycle (Pembroke et al. 2015). Single-cell transcriptomic analyses of mouse SCH neurons isolated by LCM have also revealed novel transcripts expressed in correlation with phase shifts in the circadian cycle (J. Park et al.

2016).

During shifts in circadian time, gene expression is not the only mechanism affected, but protein levels as well. Certain studies have examined proteomic changes in the whole hypothalamus after experimental disruptions in circadian rhythms (Mishra et al. 2009). Moreover, analysis of the proteome has revealed that 13% of soluble proteins expressed in the SCH undergo circadian regulation (Deery et al. 2009), and that a “time of day proteome” exists in this structure, with several proteins exhibiting marked fluctuations specifically during the transitions from light to dark and vice versa (Chiang et al. 2014). Interestingly, the SCH has become something of a model system for peptidomic studies, in that most of the peptidomic studies to date for a hypothalamic sub-region have been focused mainly on this structure (**Fig. 3D**). Peptidomic studies have revealed differential peptide abundances that correlate with changes in the time of day, including vasoactive-intestinal polypeptide (VIP) and pituitary adenylate cyclase-activating polypeptide (PACAP) (J. H. Lee et al. 2010; Southey et al. 2014). However, peptidomic signatures of the SCH do not necessarily mark peptides designated for release, and an analysis of releasates has made it possible to detect peptides designated for cell-to-cell communication (Hatcher et al. 2008; see review by Mitchell et al. 2011). Future work along these lines could help to determine differential peptide release from SCH sub-regions (e.g., the core and shell), which are known to have distinct physiological characteristics (reviewed by Moore et al. 2002). For example, neurons have a firing rhythm that need to be reset after responding to stimuli and the dynamics in gene expression patterns associated with phase resetting are different between the core and shell (Zhu et al. 2012).

#### 4.2.4: Molecular extraction from the hypothalamo-neurophypohysial system

The supraoptic nucleus of the hypothalamus (SO) is a well-studied structure known for its role in mediating fluid homeostasis and regulating parturition, and exhibits structural and functional plasticity in association with these physiological processes that signal underlying alterations in molecular expression. These hallmarks of plasticity include changes in nucleolar numbers (Hatton et al. 1972) that signify changes in ribosomal RNA synthesis; i.e., protein synthetic machinery levels (Pederson, 2011). Studies on the SO have been conducted to profile the transcriptome under normal, physiological conditions or after the effects of hypo-osmolality and/or dehydration. The main neuronal phenotypes of the SO are oxytocin (OT)- and vasopressin (VP)-expressing magnocellular neurons (MNs), which have been found to express 1,385 genes at levels that are more than twice those found in the rest of the hypothalamus, when sampled as a whole (Mutsuga et al. 2004). Taking advantage of the two types of MNs, Humerick and colleagues (2013) isolated SO MNs by their expression of OT or VP and found differential expression patterns; most notably in their transcription factors. However different these neuronal subtypes are, many studies have also examined global effects on MNs. For example, hypo-osmolality inhibits both OT and VP MNs and alters their transcriptome in comparison to the whole hypothalamus (Yue et al. 2006). Single MNs have also been isolated from rat SO and analyzed for neuropeptide phenotype markers (Xi et al. 1999; Glasgow et al. 1999; Yamashita et al. 2002; reviewed by Mutsuga and Gainer, 2006).

Together with the MNs of the paraventricular hypothalamic nucleus (PVH), the SO makes up the hypothalamo-neurohypophyseal system (HNS) that, along with several other functions, mediates fluid homeostasis. Dehydration/salt-loading can alter the HNS transcriptome, with certain genes enriched in the PVH and SO being especially sensitive to this physiological condition (Hindmarch et al. 2006; J. Qiu et al. 2011; 2014; Stewart et al. 2011; Greenwood et al. 2015; also see Hindmarch et al. 2013). Similarly, the HNS proteome is also altered by dehydration, where 25 and 45 proteins have been reported to be affected in the SO and neurointermediate lobe (NIL), respectively (Gouraud et al. 2007). K. Johnson et al. (2015) have employed next generation sequencing technology (RNA-Seq) to examine the effects of salt loading on gene expression in the SO of rats, and found that nearly 6% of the genes alter their expression levels following this intervention. Given the roles of OT and VP in the HNS system, there is also a rich interest in other peptides MNs may express. For example, Bora et al. (2008) identified 85 peptides from isolated MNs of the SO. Moreover, Hazell and colleagues (2012) provide an overview of their studies concerning the presence of various G-protein coupled receptors in the PVH and SO using high-throughput methods, along with other techniques.

Along with MNs, the PVH also harbors distinct parvicellular neurons (PNs), although their similarity is highlighted by their comparable gene expression profiles (Bonaventure et al. 2002). Of the 2,145 profiled genes within these cell types, 65% were validated via *in situ* hybridization. The PNs of the PVH that express corticotropin-releasing hormone (CRH) are involved in the stress response as part of the hypothalamic–pituitary–adrenal (HPA) axis, and distinct stressors can produce differential gene expression in the PVH (Reyes et al., 2003). Some studies on the PVH have been conducted to examine a handful of genes in PNs without technically resorting to “high-throughput methods”, such as focused studies of certain genes using real-time PCR. For example, Wang et al. (2008) examined LCM-captured human hypothalamic tissue collected *post mortem*, and identified an up-regulation of corticotropin-releasing hormone (CRH) and other gene products in associated with patients who suffered from clinical depression. Other studies have used modern “-omics” technologies to either profile the transcriptome alone (Atkins et al. 2011) or to investigate a mechanistic role for PVH genes within the HPA axis. For example, transcriptomic analysis, combined with morphometric and immunohistochemical evidence, demonstrated that select neurons, likely to be true PNs, express the gene encoding the molecule secretagogin, which is functionally linked to CRH release from these neurons (Romanov et al. 2014).

#### 4.2.5: Molecular extraction from the arcuate hypothalamic nucleus (ARH)

The arcuate hypothalamic nucleus (ARH) is a structure involved in the maintenance of energy homeostasis (see Andermann and Lowell (2017) for a recent review of ARH function within feeding control networks). Transcriptomic analyses have been conducted across multiple studies examining the effects of diet, peripheral signals, and environment on gene expression in ARH neurons. For example, Paulsen et al. (2009) identified changes in neuropeptide Y (NPY) and proopiomelanocortin (POMC) mRNAs and an additional 3,480 transcripts in fasted, diet-induced obese rats. Similarly, Jovanovic et al. (2010) showed changes in hundreds of genes in the ARH after leptin treatment in 48-hr fasted animals. Using cell sorting methods, Draper et al. (2010) isolated NPY-expressing neurons in the mouse ARH and ran microarray analysis to identify novel genes in this specific cell population in comparison to NPY-expressing neurons elsewhere in the hypothalamus (DMH), including the gene encoding the leptin receptor. At a more detailed level, Landmann et al. (2012) used LCM to sample the ARH in fasted rats, fed rats, and rats refed with a glucose load and found an up-regulation of Agouti-Related Peptide (AgRP) mRNA under fasted conditions that was greater in magnitude within single, LCM-captured neurons compared to what the authors term “ARH cell layers”, which essentially meant a complete LCM of the full ARH expanse along its cytoarchitectonic boundaries in each sampled coronal section (note the investigators performed single-cell LCM on one hemisphere, and full ARH LCM on the opposite hemisphere). In response to the refed condition, AgRP was conversely found to be down-regulated and POMC mRNA up-regulated. Importantly, the authors specified a brain atlas they used and the specific atlas levels from which they sampled the ARH, setting this study apart from most others in its more careful delineation of anatomical boundaries.

Conducting cell type-specific transcriptomics, Henry and colleagues (2015) identified molecular pathways specific to AgRP neurons that were differentially affected in fed and food-deprived animals. Similarly, Campbell et al. (2017) found using Drop-Seq methodology (see **Section 4.2.2**) that thousands of genes coding for non-neuronal and neuronal cell types displayed altered expression in association with changes in feeding conditions and energy states. They found that the transcriptional response to fasting was generally stronger than that produced by a high-fat diet, with neuronal types responsive to fasting also responsive to high-fat feeding.

Transcriptomics has also addressed questions about the relationship between the ARH and the peripheral nervous system. For example, Adler et al. (2012) characterized the transcriptome of retrogradely-labeled neurons within the ARH projecting to white adipose tissue. Neurogenin 3, a transcription factor that helps differentiate pancreatic endocrine cells also comprised a portion of the transcriptomic profile of NPY neurons of the ARH (Arai et al. 2010). Other cell types in the ARH that have been targets of molecular profiling include cholinergic neurons, many of which were found to also express tyrosine hydroxylase and markers for GABAergic neurotransmission (Jeong et al. 2016).

Transcriptomic analyses have also been used to address how the environment can affect the ARH. For example, low protein diet during postnatal development reduces body fat, and increases leptin and melanocortin receptors (Stocker et al. 2012). The ARH also maintains stability in its expression patterns under certain changes within the internal environment, such as during pregnancy. Specifically, Phillipps and colleagues (2013) showed that despite higher shifts in plasma leptin and insulin and low blood glucose induced by pregnancy, there are no changes in the ARH transcriptome. These studies have provided understanding as to what extent the ARH transcriptome is affected by environment. Finally, transcriptomics traditionally provides information about the expression levels of mRNA but can also provide valuable information expression concerning microRNA levels. A set of more than 210 microRNA genes was profiled in both the ARH and the PVH as potential regulators of mRNA (Amar et al. 2012).

In contrast to transcriptomics, only a few investigators have investigated the ARH proteome. For example, proteomic analysis of protein markers in the ARH after exposure of the organism to an inorganic compound demonstrated a few proteins that are altered in their levels of expression that are related to cell morphology, axonal growth and tissue remodeling (Amigó-Correig et al. 2012).

#### 4.2.6: Molecular extraction from other hypothalamic sub-regions (LHA, VMH)

A number of studies have performed molecular analyses of peptidergic neurons known to be enriched in the LHA, a relatively large expanse of the hypothalamus that harbors a diversity of cell types (Bonnavion et al. 2016). For example, Volgin et al. (2004) reported isolating individual slices of brain containing portions of the LHA and creating suspensions of dissociated cells from that region. They then identified the peptidergic phenotype of the cells using antibodies raised against the precursor peptide encoding hypocretin/orexin (H/O), pre-pro-H/O; or melanin-concentrating hormone (MCH); and performed RT-PCR on each cell for the respective mRNAs for these neuropeptides, providing a proof of concept for their delicate methods. Harthoorn et al. (2005) reported using single-cell LCM to generate transcriptional profiles of neurons expressing MCH and H/O, and found that these neurons express transcripts for several other neuropeptides, such as dynorphin and cocaine- and amphetamine-related transcript (CART).

Using a translational profiling technique called TRAP (Translating Ribosome Affinity Purification; Doyle et al. 2008; Heiman et al. 2008; Dougherty et al. 2010), which involves affinity purification of polysomal mRNAs in defined cell populations, Dalal et al. (2013) generated mouse transgenic lines that expressed a fusion protein encoding enhanced green fluorescent protein and the large-subunit ribosomal protein L10a (eGFP-L10a) in hypothalamic neurons that express H/O. The expression of this fusion protein allows for the isolation of those mRNAs within H/O-expressing neurons that are undergoing translation at the site of polyribosomes, effectively allowing a translational profiling of a chemically identified neuron. Using this approach, the investigators identified >6,000 transcripts with signal above background levels; 188 of these were highly enriched in H/O neurons (Dalal et al. 2013). Fifteen of these transcripts were confirmed to be present within intact H/O neurons by dual-label *in situ* hybridization, including the transcription factor Lhx9, which the authors showed, using gene ablation experiments, that it contributes to maintaining wakefulness in mice. Using an extension of the TRAP approach on the same problem, which they dubbed “vTRAP” (“viral TRAP”), Nectow et al. (2017) engineered a Cre-dependent adeno-associated virus to harbor a construct encoding eGFP-L10a, to translationally profile a specific variety of cell types in layer 5 of the cerebral cortex, the dorsal thalamus, ventral tegmental area, dorsal raphe nucleus, and LHA. Within the latter region, they focused on targeting their viral construct to MCH-expressing neurons.

The Jackson laboratory has recently reported single-cell transcriptomic data obtained from LHA H/O-expressing neurons and MCH-expressing neurons in mouse transgenic lines (Mickelsen et al. 2017). Importantly, in their study, they show specific delineations of the regions they dissected using atlas-based coordinates and drawings of the estimated areas they micropunched. A surprising finding from their careful analyses was that virtually all MCH neurons and approximately half of H/O neurons express markers for glutamate release and GABA synthesis (but not GABA vesicular release), underscoring the importance of fast-acting, small neurotransmitters within these peptidergic neurons and highlighting potentially interesting roles for GABA metabolism with glutamatergic neurons.

Studies have also been conducted to analyze the molecular expression patterns within the ventromedial hypothalamic nucleus (VMH). The Elmquist laboratory performed LCM to isolate and analyze the VMH from mice and used microarrays to detect genes enriched in this region of the hypothalamus (Segal et al. 2005). They compared the genes they obtained with those obtained from nearby regions (the ARH and dorsomedial hypothalamic nucleus; DMH). They used real-time PCR to validate nine of the twelve most robustly expressed genes, and went on to confirm the expression of three of these genes using *in situ* hybridization. Their work complements that conducted by the Ingraham laboratory, which furnished a transcriptome from manually microdissected tissue samples obtained from the developing mouse (Kurrasch et al. 2007), in which they identified and confirmed the expression of six different VMH-enriched markers from their initial screens. At the protein level, the Renner laboratory conducted studies in which they micropunched the VMH from female rats in an atlas-guided fashion, and identified several proteins that could be reproducibly resolved via 2-D gel electrophoresis from the micropunches, including several sensitive to estradiol regulation (Mo et al. 2006; 2008).

### 4.3: A note about “hypothalamic-derived” molecules

Before moving on to discuss LCM, it is worth ending this portion of the narrative with a brief note regarding molecular provenance from the perspective of evolution. In this section, we have focused on molecular extraction of molecules from the hypothalamus, including, to name a few, neuropeptides of the hypothalamo-neurohypophysial system (OT and VP), the circadian system (VIP), and wakefulness and energy balance (H/O, MCH, AgRP). However, it is important to bear in mind that these “hypothalamic-derived” molecules are not strictly linked to the vertebrate hypothalamus *per se*, since large-scale molecular phylogenetic studies have identified precursors and analogs of these molecules in animal taxa that have evolved nervous systems lacking a hypothalamus (Jékely 2013; Mirabeau and Joly 2013; E. Williams et al. 2017). For example, Semmens et al. (2016) performed transcriptomic studies of the radial nerve cords of the European starfish, *Asterias rubens*, and identified >40 neuropeptide precursors in this echinoderm, many of which have homologs in the vertebrate hypothalamus. Indeed, precursors to neuropeptides found in the mammalian hypothalamus can be found in many phylogenetically ancient animal taxa (see supplemental data of Elphick et al. 2018). Thus, in our quest to preserve the provenance of molecular data from the hypothalamic regions from which they are extracted, we must bear in mind the ironic fact that many of the molecules, from an evolutionary standpoint, never “belonged” to the hypothalamus in the first place.

## 5: Laser-capture microdissection studies: Methodological considerations

In this section, we describe how laser-capture microdissection (LCM) techniques are a useful step for precisely delineating regions of interest within the hypothalamus for subsequent high throughput molecular analyses. We describe a few approaches involving this technique and their advantages and disadvantages, followed in **Section 6** with how such samples can be traced back to their regions of extraction using digital atlas-based mapping techniques.

Since its development in the late 1990s (Emmert-Buck et al. 1996), LCM has been a useful procedure for obtaining RNA from single cells or whole regions of tissue (for selected reviews of techniques, see Espina et al. 2006; Baskin and Bastian, 2010; Datta et al. 2015). LCM has been widely used to collect individual cells (Backholer et al. 2010; J. Park et al. 2016) or groups of cells from tissue slices or cultured cells that have been identified using immunocytochemistry (termed immuno-LCM; Baskin and Bastian, 2010) or specific fluorescent tags (e.g., GFP) or fluorescent dyes such as Alexa Fluor™ 488). These approaches have enabled users to examine the expression of anywhere from a few genes of interest upwards to several hundred genes in specific cell types for various applications including genomics, transcriptomics (next-generation sequencing, microarrays; Paulsen et al. 2009), and proteomics (Moulédous et al. 2003; for a review of applications, see Datta et al. 2015). In this section, we describe findings and/or present data from our use of two different LCM systems: 1) the Arcturus AutoPix Fluorescent LCM System (Thermo Fisher Scientific, Waltham, MA) and 2) Leica LMD 7000 Microscope (Leica Microsystems Inc., Buffalo Grove, IL). In contrast to the Arcturus AutoPix LCM model in which the dissected tissue was collected onto a plastic cap (CapSure LCM Caps) *above the slide*, both dissected tissue and membrane surrounding the tissue was collected *below the slide* containing a UV-absorbing membrane (MembraneSlide) into a microcentrifuge tube cap using the Leica LMD7000 Microscope. Both LCM instruments have now been replaced by more recent models, including the ArcturusXT™ LCM System (now distributed through Thermo Fisher Scientific) and the Leica LMD6/LMD7. Here, we present ways in which LCM has been used to collect: (1) regions of tissue from anatomically distinct areas of the brain (**Section 5.1**); and (2) targeted populations of cells that have been identified using immunocytochemistry (**Section 5.2**) or fluorescent conjugates (**Section 5.3**). We discuss advantages and pitfalls to using these approaches.

### 5.1: LCMfor general sampling of brain regions

This is the most common application of LCM for collection of brain tissue involves collecting anatomically matched regions of tissue across several rostrocaudal levels of a particular brain site. For example, we have used the Arcturus AutoPix Fluorescent LCM System to confirm sufficient knockdown of OT receptor mRNA following hindbrain nucleus of the solitary tract (NTS) injection of OT-saporin toxin relative to control saporin toxin (Baskin et al. 2010). We collected bilateral samples of NTS tissue from slide-mounted cryostat sections (10 μm) at the level of the area postrema (AP) and rostral to the AP at 200 μm intervals (n=8 slides/brain). Following LCM collection, sections were dehydrated in ethanol and xylene, and then air-dried. We have found that this approach was suitable for measuring differences in NTS expression of OT receptor mRNA. In addition, we have used the Arcturus AutoPix Fluorescent LCM System to confirm the “expected” reduction in cholecystokinin-1 receptor (CCK1R) mRNA in both the ARH (−3.48 mm to −2.04 mm from Bregma; Paxinos and Watson 2007) and dorsomedial hypothalamic nucleus (DMH) (−3.60 mm to −2.80 mm from Bregma; Paxinos and Watson 2007) in rats that lack CCK1Rs relative to wild-type rats (Blevins et al. 2012). As before, slide-mounted cryostat sections (10 μm) of ARH and DMH were selected at 200 μm intervals, dehydrated in ethanol and xylene, and then air-dried (n=6 slides/brain). Bilateral samples were collected from brain sites that normally express CCK1R (i.e., ARH and DMH). Lastly, we have used the Arcturus AutoPix Fluorescent LCM System (**Figure 4**, *left panel*) and Leica LMD 7000 Microscope (**Figure 4**, *right panel*) to confirm the increase of NPY/ AgRP in the ARH from 48-h fasted rats relative to *ad libitum* fed rats (T. Hahn et al. 1998; Mizuno and Mobbs 1999; Korner et al. 2001; Swart et al. 2002; Bi et al. 2003). We collected bilateral samples of ARH (−3.48 mm to −2.04 mm from Bregma; Paxinos and Watson 2007) from slide-mounted cryostat sections (10 μm) at 200 μm intervals (n=6 slides/brain). In all cases, sections from adjacent slides were stained with cresyl violet (Blevins et al. 2000a,b; Baskin et al. 2010) to enable anatomical matching. As noted earlier, Panels B and C of **Figure 2** show an example of LCM of the ARH from cresyl violet-stained tissue sections. Landmann et al. (2012) have extended these findings by using LCM to demonstrate that fasting results in increased AgRP mRNA expression from the ARH (both when collected as a region or as single neuron pools consisting of 100 neurons). LCM has been used by other labs to profile the molecular composition of various hypothalamic regions (**Tables 1–3**). For example, to highlight a few studies by way of illustration, LCM has been used to confirm: 1) the effectiveness of adeno-associated viral knockdown of angiotensin II receptor subtype 1a in the subfornical organ (SFO) of rat brain (Walch et al. 2014); 2) reductions in gene expression in brains of steroidogenic factor 1 (SF-1) in the VMH of knock-out mice (Segal et al. 2005); and 3) fasting-elicited changes in gene expression in the PVH and the impact of leptin replacement on these genes (Tung et al. 2008).

**Fig. 4.**
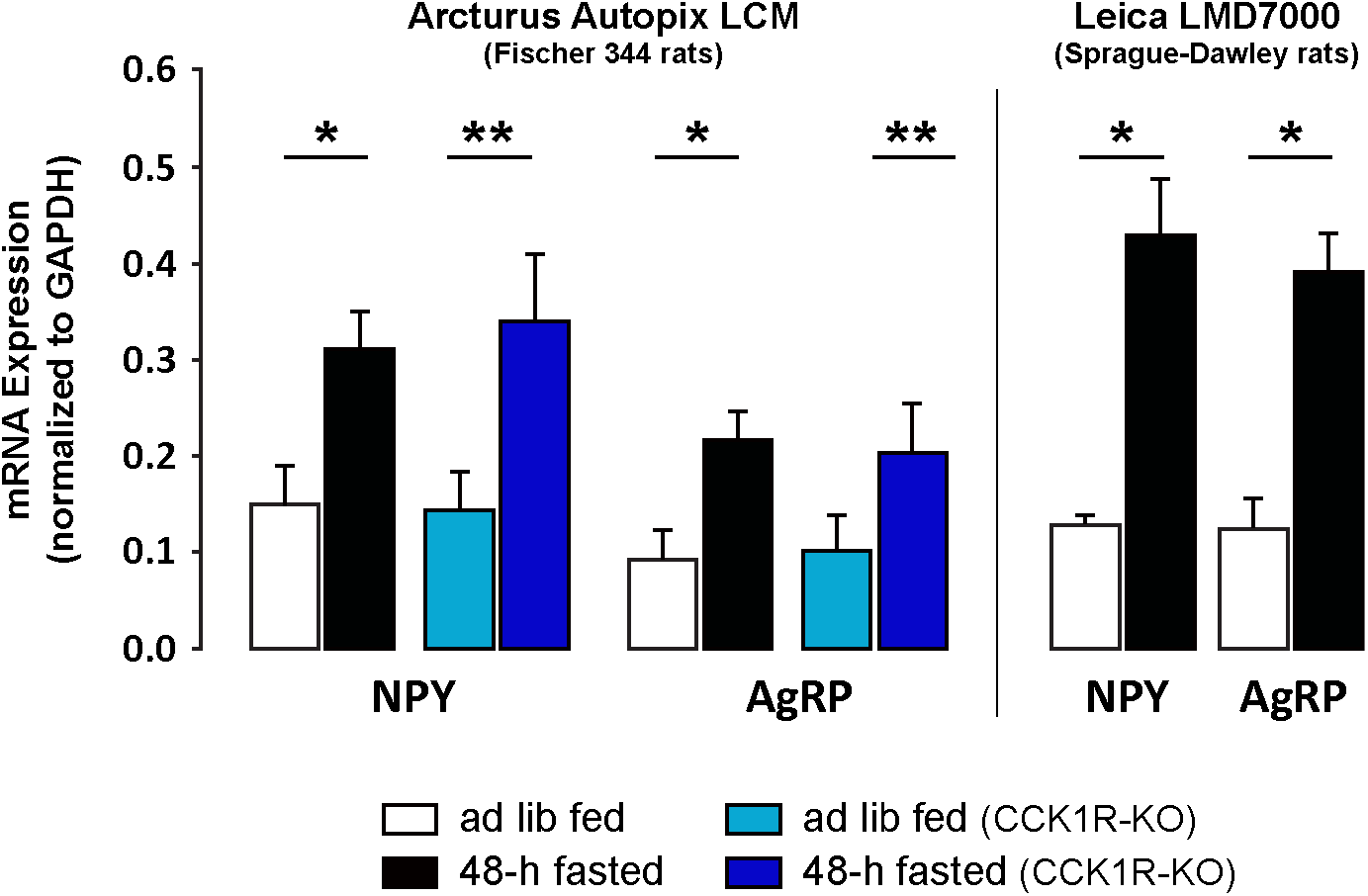
Unpublished data furnished here to illustrate the efficacy of two LCM systems (Arcturus and Leica), along with qPCR, to confirm fasting-elicited increases in NPY and AgRP mRNA expression in the ARH in rats. Slide-mounted cryostat sections (10 μm-thick) were generated through the ARH from *ad libitum* fed or 48-h fasted wild-type or cholecystokinin-1 receptor (CCK1R) knockout (KO) Fischer 344 rats (**left panel**) or *ad libitum* fed or 48-h fasted wild-type Sprague-Dawley rats (**right panel**). Sections were thawed briefly prior to dehydration in ethanol and xylene and allowed to air dry as described previously (Blevins et al. 2000a; Baskin et al. 2010; Blevins et al. 2012). LCM was used to collect bilateral sections (10 μm-thick) from the ARH (200 μm intervals) from six anatomically matched levels and sections were mounted onto slides. All microdissected samples from a single brain were pooled for RNA extraction. NPY and AgRP mRNA expression were measured using qPCR. Data are expressed as mean ± SEM. \**P*<0.05 or \*\**P*=0.1 fed vs. fasted.

### 5.2: Immuno-LCM

#### 5.2.1: Advantages

Immuno-LCM (Fassunke et al. 2004; Fink et al. 2000; Waller et al. 2012) is the approach of using immunocytochemistry to identify cells to be collected by LCM. One of the primary advantages of immuno-LCM is that it enables the user to phenotype specific cells of interest that could not be as easily identified using anatomical landmarks alone. This may be a particularly useful approach given that tissue sections collected by LCM cannot be coverslipped and, as a result, may not allow sufficient resolution to identify anatomical landmarks readily. One of our laboratories (JEB) has used this approach to identify (following a rapid immunostaining procedure for tyrosine hydroxylase (TH); a marker of catecholamine neurons) and collect catecholamine immunopositive neurons from the A2/C2 catecholamine cell groups in the hindbrain NTS. We used the Arcturus AutoPix Fluorescent LCM System to confirm the specificity of this approach by measuring TH mRNA from TH+ neurons relative to TH− neurons in order to confirm its presence for qPCR analysis (D. Williams et al. 2008). There are a number of protocols for rapid immunostaining that have already been published (Fink et al. 2000; Uz et al. 2005; D. Williams et al. 2008; Baskin and Bastian 2010; Carreño et al. 2011; Nedungadi and Cunningham 2014; J. Park et al. 2016).

#### 5.2.2: Challenges and Pitfalls

There are a number of challenges when using the rapid immunostaining approach that must be considered prior to incorporating immuno-LCM. For example, as reviewed by Baskin and Bastian (2010), the process of immunostaining can introduce the potential for RNA extraction and degradation. In an effort to minimize loss and degradation of RNA, common strategies are to implement rapid immunostaining protocols and the use of alcohol fixation (methanol or ethanol) in place of formaldehyde-based fixatives (which can result in much of the RNA being fragmented and degraded by formalin) (Gillespie et al. 2002; Soukup et al. 2003; J. Su et al. 2004; Hu et al. 2005; Baskin and Bastian 2010; Carreño et al. 2011; Nedungadi and Cunningham 2014; J. Park et al. 2016). We have previously shown that brief thawing (~30–60 sec) of cryostat-cut sections of frozen rat brain in combination with quick immunostaining after methanol fixation (~3 min) works well for immuno-LCM and qPCR for mRNA (D. Williams et al. 2008). Other challenges to the use of the rapid immunostaining approach include antibodies that require a low titer or are relatively nonspecific as well as antigens that are found in low-abundance (Baskin and Bastian, 2010). As Baskin and Bastian (2010) indicate, adjustments in staining times, incubation temperatures or more sensitive fluorochromes, may increase the specificity to acceptable levels. Rapid immunostaining approaches may be less suitable for targeting and collection of cells with low gene expression.

Another challenge when selecting specific cells is that contamination from neighboring cells may also be included in the sample. For example, Okaty et al. (2011) reported in their metaanalysis of various cell isolation methods conducted by certain laboratories that LCM produced higher contamination from spurious signals, as compared to other cell isolation methods, such as TRAP, FACS, immunopanning, and manual sorting of fluorescently labeled cells. Their analysis included an immuno-LCM study (C.-Y. Chung et al. 2005) and one in which LCM was performed on fresh-frozen brain tissue sections containing genetically labeled cells from transgenic mice (Rossner et al. 2006). One means to address this issue is to collect an equal number of neighboring cells outside of the intended region of analysis as negative controls to run alongside the positively labeled cells. We have found that selecting ~150-200 TH+ and adjacent non-catecholaminergic cells (TH−) cells from several adjacent sections was a suitable approach for measuring increases in TH mRNA from TH+ cells relative to TH− cells from the A2/C2 catecholamine cell groups in the hindbrain NTS (D. Williams et al. 2008).

### 5.3: Use of LCM to target cells expressing fluorescent reporter molecules

#### 5.3.1: Advantages

Similar to immuno-LCM, this approach enables the phenotyping of specific cells of interest that could not be as readily identified using anatomical landmarks. In contrast to immuno-LCM, there is no need for rapid immunostaining as the fluorescent tag is already present. We have used this approach previously (Blevins et al. 2009; see **Figure 2D–F**) to identify those neurons in the PVH that project to the hindbrain NTS using Alexa Fluor™ 488-conjugated retrograde tracer, cholera toxin subunit B (CTB). We have found that brief thawing (~30–60 sec) of cryostat-cut sections of frozen rat brain, in combination with selecting ~250 CTB+ cells from three or four anatomically matched coronal sections from PVH, was a suitable approach for measuring OT, CRH, and melanocortin-4 (MC4-R) receptor mRNAs (Blevins et al. 2009). We also collected the same number of neurons from the SCH as a negative control, as this site expresses relatively low levels of each of these transcripts (Mountjoy et al. 1994; Jing et al. 1998). In addition, unlabeled cells from the PVH were collected and screened for OT mRNA, CRH, and MC4-R mRNAs.

#### 5.3.2: Challenges and Pitfalls

One potential limitation of using LCM to collect GFP-labeled cells is that free GFP is soluble and can leak out from cryostat-cut sections in the absence of fixation (Jockusch et al. 2003), thus necessitating perfusion and/or post-fixation of the tissue. Soluble eGFP is preserved in paraformaldehyde (PFA)-fixed tissues that are post-fixed in 50% ethanol and 100% *n*-butanol (Khodosevich et al. 2007). The authors noted that while PFA fixation of mouse tissue is sufficient in preserving the EGFP signal for up to 30–60 min, it was not sufficient in preserving EGFP signal for longer periods of time (Khodosevich et al. 2007). They also indicated that post-fixation in alcohol is “necessary not only to remove the water to prevent RNA degradation, but also to render the aldehyde-crosslinks more stable, thus preserving the fluorescence” (p. 2). They added that “alcohol fixation alone also was not sufficient to preserve fluorescence of the soluble EGFP and prevent it from leaching out and diffusing to neighboring tissue making it impossible to specifically identify green fluorescent cells” (p. 2). Although some groups have reported relative disadvantages of using formaldehyde-based fixatives to retrieve PCR product from LCM-sampled non-neural (Goldworthy et al. 1999) and neural tissues (J. Su et al. 2004), there are instances where LCM has been shown to work successfully on formaldehyde-fixed tissues (e.g., Kabra et al. 2016). Recent papers indicate that EGFP+ (or EYFP+) cells can also be harvested from fresh frozen mouse (Rossner et al. 2006; Liu et al. 2011) and rat brain tissue (Liberini et al. 2016), but the extent to which the fluorescent signal may have diffused or faded beyond 30–60 min were not addressed in these studies. It is worth noting that Leica has produced a protocol designed to optimize visualization of GFP from post-fixed tissue to be used for LCM.

### 5.4: RNA Integrity

The RNA Integrity Number (RIN) value is a tool developed by Agilent Technologies to assess RNA integrity using the Agilent 2100 Bioanalyzer and RNA LabChip^®^ kits. The RNA integrity is based on the electrophoretic trace of the sample and allows the user to assess the amount of degradation products in the sample and to determine integrity of the sample. It is an important consideration when assessing gene expression data from samples generated by LCM. The RIN algorithm assesses RIN values that range from 1–10 with 1 representing completely degraded RNA, 5 representing partially degraded RNA, and 10 representing completely intact RNA. We have used the 2100 Electrophoresis Bioanalyzer (Agilent Technologies) to obtain RIN values from ARH tissue samples that had been stored for ~3 months at −80°C. We obtained RIN values ranging from 7.6–8.2 (7.92 ± 0.09). These RIN values are comparable to those we obtained from ARH tissue (7.8 and 8.5) that had been stored ~7–8 months at −80°C. While these values are in the higher range it does indicate some degree of degradation. These findings are also consistent to the RIN values (6.2) reported from tissue collected from patients with oral cancer that was stored for ~48 h at −80°C (Xu et al. 2008), as well as RIN values (6–7) reported from pancreatic tissue collected from rats and humans (Butler et al. 2016). They are also consistent with RIN values (6.6–7.6) reported for hypothalamic tissue sampled using LCM from the supraoptic (SO) nucleus; the LCM was performed within one month following tissue sectioning and storage of the slide-mounted sections at −80°C (K. Johnson et al. 2015).

## 6: Anchoring molecular information to their native regions using digital atlas maps

Having reviewed in the preceding sections the importance of location information in the brain (**Section 2**), the historical antecedents of current high throughput work concerning molecular extraction of the brain (**Section 3**) and the hypothalamus (**Section 4**), and the methodology of LCM (**Section 5**); we now turn to the topic that constitutes the principal thesis of this review; namely, the mapping of datasets to standardized atlases of the brain. Using the backdrop of LCM procedures described in the preceding section, we discuss first how documenting the location of the native substrate from where tissue is extracted is critical for the subsequent mapping of that location, and then describe the mapping steps themselves.

### 6.1: Documenting the native substrate before extraction

Applying LCM to a tissue section to capture and sample a particular region of interest can be performed in a number of ways, a few of which were described in **Section 5**. Unstained tissue sections can be viewed under a dark field microscope to observe the region of interest in relation to white matter tracts that might be nearby. Such landmarks can aid greatly in the accurate and repeated sampling of a region, especially for large sub-regions of the hypothalamus, a part of the brain replete with white matter landmarks (e.g., anterior commissure, optic chiasm, optic tract, fornix, mammillothalamic tract). Indeed, what is perhaps the first documented sampling of the hypothalamus was reported diagrammatically in relation to many of these fiber systems (see **Fig. 2A**). Micropunch methods, first developed before the establishment of LCM, involve procedures where tissue punches are harvested from unstained frozen or fresh tissue sections; in such cases, white matter tracts also serve as important landmarks to orient the experimentalist as to where a particular region of interest was located and how much tissue to collect from that region (Palkovits 1973; Jacobowitz 1974).

Apart from unstained tissue, the most common method for identifying regions of interest in sectioned brain tissue is through the use of Nissl stains (stains that label basophilic substrates – ‘Nissl substance’ – in the cell, including rough endoplasmic reticulum and the nucleus, sites where nucleic acid molecules are concentrated). The use of such stains on brain tissue sections prior to LCM-based sampling from those sections is a common way of accurately delineating regions of interest for LCM-targeting (e.g., Ginsberg and Che, 2004; Boone et al. 2013). Nissl-based stains such as cresyl violet (**Fig. 2B**), thionin, and hematoxylin have been used to guide sampling of hypothalamic sub-regions and cells, including the preoptic region (Vasilache et al. 2007; Aubert et al. 2013), ARH (Segal et al. 2005; Jovanovic et al. 2010), SCH (Porterfield et al. 2007; Boone et al. 2013; Pembroke et al. 2015), SO (Xi et al. 1999; Glasgow et al. 1999; Yamashita et al. 2002; S. Wang et al. 2008), VMH (Segal et al. 2005; Kurrasch et al. 2007), DMH (Segal et al. 2005), and PVH (S. Wang et al. 2008; Blevins et al. 2009; Novoselova et al. 2016). Importantly, investigators have performed LCM on the Nissl-stained tissue itself (Segal et al. 2005; Porterfield et al. 2007; Jovanovic et al. 2010; Boone et al. 2013), but in principle, one can also use adjacent sections stained for Nissl substance to help delineate regions of interest on unstained companion sections sampled by LCM, as has been done for human tissue samples collected *post mortem* for the PVH and SO (S. Wang et al. 2008). In addition to using Nissl staining as a tool to help delineate LCM-captured tissue sample boundary conditions, other stains and labeling strategies have also been used in conjunction with LCM, including FluoroJade for delimiting tissue pathology (Boone et al. 2013), Cy3-conjugated secondary antibody to identify antibody-labeled peptidergic neurons (Nedungadi et al. 2012a,b; Nedungadi and Cunningham 2014), immunoperoxidase-based detection of peptidergic neurons (Briski et al. 2010), NeuroTrace staining for visualizing fluorescent Nissl-like profiles (Benner et al. 2015), and *in situ* hybridization in human *post mortem* tissue (Bernard et al. 2009). Finally, it is worth noting that although LCM procedures themselves do not appear to result in significant losses of protein as compared to manually dissected samples of comparably located regions, Nissl staining itself can be detrimental to the full retention of some proteins for subsequent proteomic analyses (Moulédous et al. 2002), and the use of Nissl stains such as neutral red, cresyl violet, or NeuroTrace reportedly contributes to lower yields of transcripts from LCM-captured brain tissue (Kerman et al. 2006; Benner et al. 2015).

### 6.2: Mapping to standardized atlases

Using aids such as the Nissl stain to identify a region of interest to be sampled by LCM not only helps ensure accurate sampling of that region, but also provides an opportunity to document the location of the excised tissue itself using standardized atlases of the brain. Such atlases have existed for several decades, and many have been created for a variety of animal models, including – to name but a few – toads (Hoffmann 1973), frogs (Wada et al. 1980), lizards (Greenberg 1982), guinea pigs (Tindal 1965), rabbits (Sawyer et al. 1954), mice (Dong 2008; Paxinos and Franklin 2012), and rats (Swanson 2004, 2018; Paxinos and Watson 2007) (for a detailed listing, see Ten Donkelaar and Nicholson 1998). As detailed in Khan (2013), there are many advantages of using standardized atlases to map experimental data, not least of which is to be able to spatially align different datasets from diverse studies and contextualize them with some rigor and precision (also see Khan et al. 2018).

How is mapping experimental data to a reference atlas of the brain performed? Simmons and Swanson (2009) describe many aspects of how mapping experimental data to a standardized reference atlas is undertaken. A critical factor is reconciling the plane of section of the experimental tissue with the plane of the atlas map that will be used to contain the mapped dataset. Differences in plane of section, determined by the angle of cutting on the microtome or cryostat instrument used to section blocks of brain tissue, can potentially constitute a significant source of mapping errors, especially in the absence of any global (e.g., Nissl) stain to mark the cytoarchitecture of the tissue being sectioned. It is surprising to us how few investigators explicitly discuss how they have dealt with plane of section issues when analyzing the results of expression and distribution studies.

For example, many investigators have utilized immunohistochemistry of the transcription factor and immediate-early gene product, Fos, to identify regions of the brain *post mortem* that may be associated with patterns of activation or with particular behaviors that the organism was involved in during the life history immediately preceding death. However, to our knowledge, none of these studies presents a comparison of Fos expression patterns between groups of animals while providing an explicit discussion of how planes of section were taken into account in their determination of regional comparisons. Thus, as part of a collaborative study (Zséli et al. 2016), a few of us (AM, AMK) performed a plane of section analysis to map patterns of Fos expression in rats who had fasted for 40 h (but had *ad libitum* access to water) versus rats who fasted for 40 h but then were allowed to re-feed for 2 h. **Figure 5** shows a portion of these data. Compared to the plane of section of the Swanson (2004) reference atlas (**Fig. 5A**, *top panel*), the planes of section for the subjects examined for Fos expression were markedly different (**Fig. 5A**, *middle and bottom panels*), and any accurate comparison of the same regions at similar rostrocaudal levels between fasted and re-fed cohorts required reconciling the planes of section for tissues sectioned from both cohorts with the plane of section of the reference atlas. This was not only important for representing the patterns of expression on the atlas, but also to ensure that we were not comparing levels of expression between regions that did not correspond with one another in terms of spatial positioning within the brain.

**Fig. 5.**
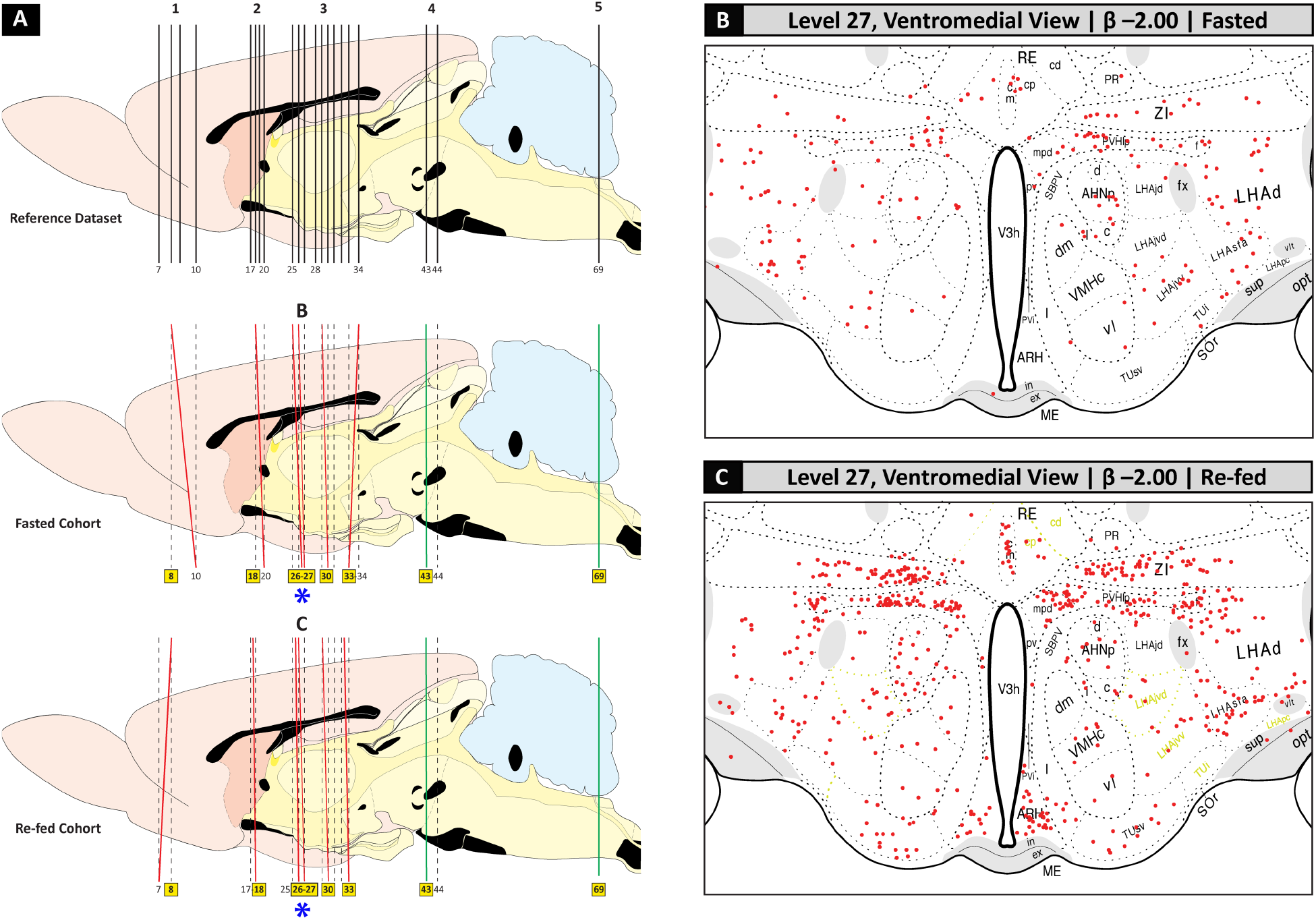
Example of a plane-of-section analysis conducted on Fos transcription factor expression maps of hypothalamic tissue sections obtained from fasted *versus* refed adult male Wistar rats, conducted by the UTEP Systems Neuroscience Laboratory as part of a published collaborative study with the Hungarian Academy of Sciences and Tufts University (Zséli et al. 2016). **(A)**. Sagittal views of the rat brain, modified from Swanson (2004), showing the plane of section of the Swanson atlas (*top panel*: ‘Reference Datase’), followed by the planes of section for a subject from the fasted cohort (*middle panel*) and the refed cohort (*bottom panel*). **(B, C)**. Locations of Fos-immunoreactive neurons plotted onto coronal-plane maps from Swanson (2004) showing the ventromedial views of atlas level 27 for each cohort (which are denoted by *asterisks* in **A**). Reproduced from Zséli et al. (2016) and with permission from John Wiley & Sons, Ltd.

As detailed in our study (Zséli et al. 2016), we utilized the digital atlas maps for Swanson, (2004), which are now also available online (https://larrywswanson.com). These were manipulated in Adobe Illustrator (AI) software. Nissl counterstain in the Fos-immunoreacted tissue sections was used as a guide to identify cytoarchitectural boundaries for each section. The photomicrographs were imported into separate layers of AI, scaled, and compared to the atlas plates to determine whether there were differences in plane across the mediolateral and dorsoventral axes. In some cases, as delineated by Simmons and Swanson (2009), patterns on a tissue section require mapping to more than one level of a reference atlas, and the differences in plane of section are often in more than one plane simultaneously, necessitating a segment-by-segment translation of the region of interest to the relevant location on a map or set of maps.

Another important point to note about deriving information for mapping on the basis of Nissl-stained tissues is that often Nissl stains do not fully reveal distinct patterns of cytoarchitecture within tissue; in such cases, it is at times difficult to discern a particular sub-region within a tissue section and determine precisely the boundaries of a region. In such cases, we have opted to report the uncertainty in our mapping that results from such ambiguous staining patterns, by noting within the reference atlas those portions of the map that are based on inferred positions of cytoarchitectonic boundaries as opposed to those that were directly observed (and distinct) within the stained tissue section. As shown in **Figure 5C**, we found certain sub-regions of the LHA to display Nissl patterns that were indistinct, which permitted us to only infer positions of the Fos-immunoreactive cells we were mapping. This uncertainty was represented in the form of a pale yellow color for the dotted line boundaries for those regions (**Figure 5C**).

For LCM-captured brain tissue, an outline of basic steps for mapping the sampled tissue to a reference atlas is shown in **Figure 6**. First, as described in **Section 5.2.2** and **Section 5.3.2**, investigators have to decide whether to employ fixatives such as methanol, alcohol or formaldehyde to preserve their tissues of interest before sectioning them, or instead opt to use freshly frozen, unfixed tissue sections (**Fig. 6A**, *Step 1*). Once sectioned and mounted onto slides (**Fig. 6A**, *Step 2*), a given tissue series can be Nissl-stained (**Fig. 6A**, *Step 3*), and then placed within an LCM instrument to excise a region of interest (ROI; **Fig. 6A**, *Step 4*). Apart from the sequestration and processing of the LCM-captured tissue of interest for further analyses using transcriptomics, proteomics, or peptidomics, etc., the remaining tissue section (i.e., the rest of the section that remains *after* the region of interest has been excised) can now be used as a key to unlock the precise location of the sampled area within a standardized reference space of the brain. Similar to the example of a plane-of-section analysis furnished in **Figure 5**, the section can be examined in relation to the Nissl-based landmarks of photographs within the reference atlas to be used, and the tissue’s plane of section assigned to appropriate levels of the reference atlas (**Fig. 6B**, *Step 5*). The ROI within the tissue section can then be mapped using a digital atlas map of that reference level.

**Fig. 6.**
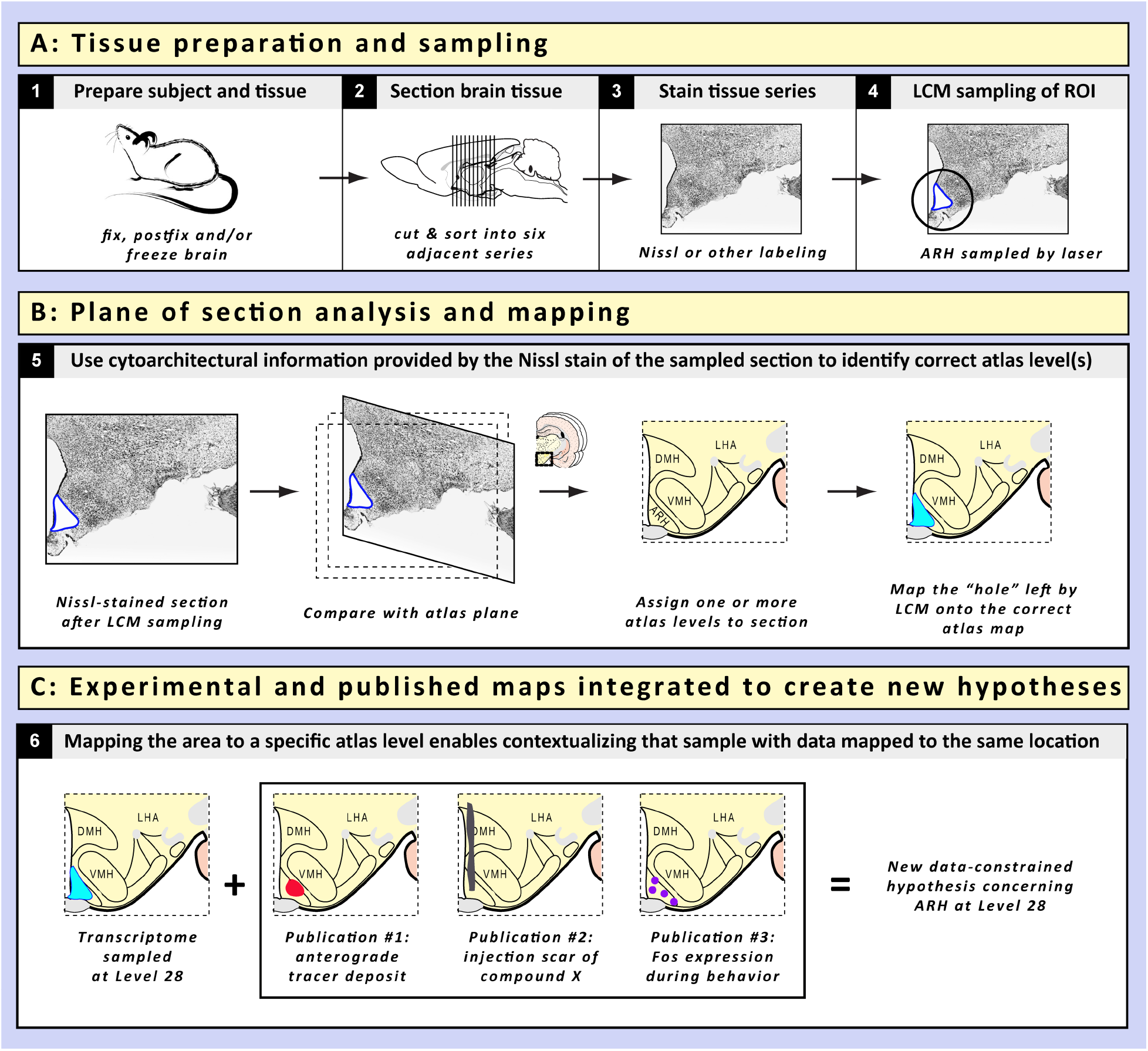
Steps for mapping tissue samples collected using LCM, and the advantages of such mapping. Following steps to process brain tissue through the tissue fixation (**Panel A**, *Step 1*) and sectioning/staining/LCM-based sampling steps (**Panel A**, *Steps 2–4*), the remaining mounted and Nissl-stained tissue section from which the sample was excised can be mapped and analyzed for the location of the ‘hole’ left from sampling in relation to a standardized reference atlas of the brain (**Panel B**, *Step 5*). The hypothetical example shown here is for LCM-based capture of the arcuate hypothalamic nucleus (ARH) (**Panel A**, *Step 4, circled region* on the Nissl photomicrograph; the white ROI outlined in *dark blue* shows the region that was ‘sampled’). **Panel C**: Once mapped, the location of the sampled tissue can be examined in relation to any other dataset that has been mapped to the same atlas level (*Step 6*). In this example, three hypothetical published datasets (a tract-tracing deposit, a drug injection site, and the distribution of activated immediate-early gene products) have all been mapped from different brains to the same reference location, thereby allowing a contextualization of the data in a spatially rigorous manner to generate new hypotheses concerning the ARH at that rostrocaudal level.

## 7: The benefits of mapping native substrates and anchoring datasets

### 7.1: Data integration

**Figure 6C** provides a view of the types of benefits that can be obtained by assiduously mapping a tissue sample obtained by LCM to a reference map of the brain. In addition to the data generated from the high throughput “-omics”-based extraction and analysis of the sample itself, the precise mapping of the sample in relation to its native landscape allows one to examine all previous studies that have been conducted on that sampled region that have been mapped within the same reference space. For example, for the Swanson (2004) reference atlas of the rat brain, several studies have utilized the digital maps of this work to map the datasets from their studies of the hypothalamus. Such studies include those involving central microinjections of molecules into the PVH (Khan et al. 2007; reviewed in Khan et al. 2017), protein expression in the LHA in response to water deprivation (S. Yao et al. 2005), transcription factor activation in several hypothalamic regions in response to fasting or re-feeding (Zséli et al. 2016), deposits of neuroanatomical tract tracer molecules into any of several hypothalamic regions (e.g., Hahn and Swanson, 2010), and mapping of key neuropeptides within distinct subdivisions of the LHA (Swanson et al. 2005; J. Hahn 2010).

In the hypothetical scenario furnished in **Figure 6C**, the location of the portion of the ARH sampled by LCM maps to Level 28 of Swanson (2004). Specifically, this region – at this same rostrocaudal level – could also have been the focus of investigations concerning anterograde tracing, central drug injection, and Fos transcription factor expression. Therefore, all of those published datasets could be considered in conjunction with the molecular analyses performed on the LCM-sampled ARH, and new hypotheses can be constructed that are constrained by the spatial patterns of data from these maps, when they are considered as a collective (**Figure 6C**). Together, therefore, the maps constitute a powerful way to help investigators see relationships among datasets, for the same region mapped in the same reference space, that they otherwise may not have seen or which they may have seen without any rigorous constraint placed upon such examination. The gene expression changes observed in molecular analyses of the sampled region, for example, may be occurring in neurons in that region for which anterograde tracing experiments have revealed prominent efferent connections. Thus, linking the molecular with neuroanatomical data would suggest new experiments that could test whether those genes play a role in shaping the function of those projection neurons. Setting aside hypothetical scenarios for a moment, we have recently reported the usefulness of this approach in a preliminary examination of published datasets for the LHA; specifically, those studies that have been performed that report LHA datasets mapped in Swanson reference space (Hernandez and Khan 2016).

### 7.2: Data migration

A logical extension of contextualizing datasets mapped to the same reference space would be to migrate data from a different reference space to the reference space one has used to map the location of their LCM-sampled tissue. Thus, as in the example furnished earlier, if the ARH sample captured by LCM were mapped to Level 28 of Swanson, it would be interesting to determine whether data concerning this region, but which was mapped in a different atlas, could be “migrated” to this reference space and contextualized with the data obtained from the LCM sample at the same atlas level. This has been discussed in detail by one of us previously (Khan 2013) and the details are not necessary to enumerate here again; suffice it to say that registration of data between atlas spaces – when performed under careful, lawful parameters – allows researchers to unlock the potential of data that may be residing, unattended, in a different reference space. This is important because many researchers use different atlases to map their datasets; this is true for the hypothalamus as much as any other brain region. For example, the locations of recording electrodes used to perform electrophysiological recordings of neurons in the PVH have been mapped to the atlas of Paxinos and Watson, along with inferred stereotaxic coordinates for the locations of the maps (Aramakis et al. 1996). The recordings are for responses these PVH neurons have to application of NPY or its receptor agonists, and understanding the locations of the neurons displaying these responses could be better contextualized in relation to other datasets mapped in Swanson reference space if the data were migrated to that space.

Fortunately, the alignment and registration between these atlas spaces appear to constitute a tractable problem (Wells and Khan 2013; Khan 2013; Hernandez and Khan 2016; Perez et al. 2017), the mature, fully fledged solution for which may help to bring together datasets that would otherwise be separated in time and space. As a step towards such a solution, we have recently developed and implemented a computer vision algorithm that matches features detected in photomicrographs of the Nissl-stained sections of the Paxinos and Watson and Swanson reference atlases to provide independent support of alignments we performed separately between the reference atlases based on craniometric measures in relation to the skull landmark, Bregma (Khan et al. 2018). The algorithm produces matches between atlas levels that are in close agreement with matches produced on the basis of craniometric alignments, providing support for the feasibility of data migration between the two reference spaces.

Other, older datasets could also be potentially migrated between atlas reference spaces, provided that the reference spaces can be aligned and registered in a fashion similar to that described above for the Paxinos and Watson / Swanson reference atlases. For example, Jacobowitz and colleagues combined micropunch methods with two-dimensional gel electrophoretic separation methods to generate protein profiles from discrete sub-regions of the hypothalamus and other brain regions (Heydorn et al. 1983), mapping their data using coordinates derived from the König and Klippel (1963) rat brain atlas. In principle, such data can be contextualized more broadly if they were migrated to other extant reference spaces.

### 7.3: Data refinement

Another benefit of mapping the location of LCM-captured brain tissue is the ability to improve our understanding of hypothalamic organization by refining the data generated from previously published studies. Prior to the advent of LCM (Emmert-Buck et al. 1996), the ability to sample brain tissue with high spatial resolution found perhaps its most precise expression in the micropunch methods mentioned above (reviewed by Palkovits 1975; 1986; 1989). Notwithstanding notable examples using these and other methods (e.g., Heydorn et al. 1983), LCM offers investigators the ability for an even greater precision of sampling of brain tissue within a given region’s 3-D expanse, thereby allowing more careful examination of sub-regions to detect possible differences in molecular expression patterns within a defined neural substrate.

This level of spatial resolution is important, as data has emerged that suggest heterogeneous neuronal constituents along the rostrocaudal extent of hypothalamic nuclei and areas. For example, within the ARH, data from the mouse model demonstrate a segregation of the effects of acutely administered leptin and insulin on populations of ARH POMC neurons (K. Williams et al. 2010). Specifically leptin-induced excitation was found in 35% of all POMC neurons throughout the rostrocaudal extent of the retrochiasmatic area (RCH) and ARH, but most of the POMC neuronal excitation was recorded from neurons in the lateral RCH and medial POMC group in the ARH. In contrast, insulin-induced hyperpolarization of POMC neurons was restricted to medial RCH and rostromedial ARH (K. Williams et al. 2010). More recently, Lam et al. (2017) used single-cell RNA sequencing to determine that the POMC neuronal population in the mouse ARH consists of heterogeneous populations that differ on the basis of their cell surface receptor expression. Clustering analysis resulted in the investigators identifying four different classes of POMC neuron. Similarly, an elegant study by Foster et al. (2016) has demonstrated the presence of distinct subsets of neurons in the VMH in the rat model that show a selective *absence* of Fos immunoreactivity in association with the hypoglycemia produced by systemic insulin injections. In particular, they found that the VMHdm (dorsomedial part of the VMH) and the smaller VMHc (central part) show marked reductions in Fos immunoreactive neurons from hypoglycemic animals as compared to their euglycemic controls, and that these reductions were proportional to the reductions in terminal plasma glucose concentrations. In contrast, sub-regions such as the VMHvl (ventrolateral part), which are believed to be involved mainly in social and reproductive behaviors, do not exhibit such reductions. Clearly, then, sampling from these smaller sub-regions of the ARH and VMH warrants careful documentation and mapping.

Our own preliminary data (Martinez et al. 2015; 2016) on ARH connectivity underscores this point as well. Specifically, initial experiments in which the retrogradely transportable tracers, Fluorogold or CTb, were injected into the rostral and caudal portions of the ARH have yielded results showing subtle differences in the distribution and density of retrogradely labeled neurons throughout the forebrain that project to these portions of the ARH. A summary of these unpublished data is furnished in **Figure 7**, simply to emphasize the point that it is no longer tenable to sample only one tissue section of a large expansive brain region such as the ARH, and make claims about its function as a whole without taking into consideration the possibility of heterogeneous properties for neurons along its full extent. A difference in afferent input implies different qualities for incoming signals to ARH neurons in the rostral end of the structure versus its caudal end; this in turn, implies that perhaps the neuronal populations receiving these differential signals may also be heterogeneous. Therefore, their molecular expression patterns, in terms of either phenotype or intrinsic state (or both) will likely also be non-uniform. Moreover, sex-specific differences in gene expression have also been reported for the ARH (Mozhui et al. 2012). Thus, the greater spatial resolution afforded by LCM sampling methods – together with careful digital atlas mapping of those locations of those samples – allows us as a community to continually refine our coarse datasets, rendering them sharper and more information-rich.

**Fig. 7.**
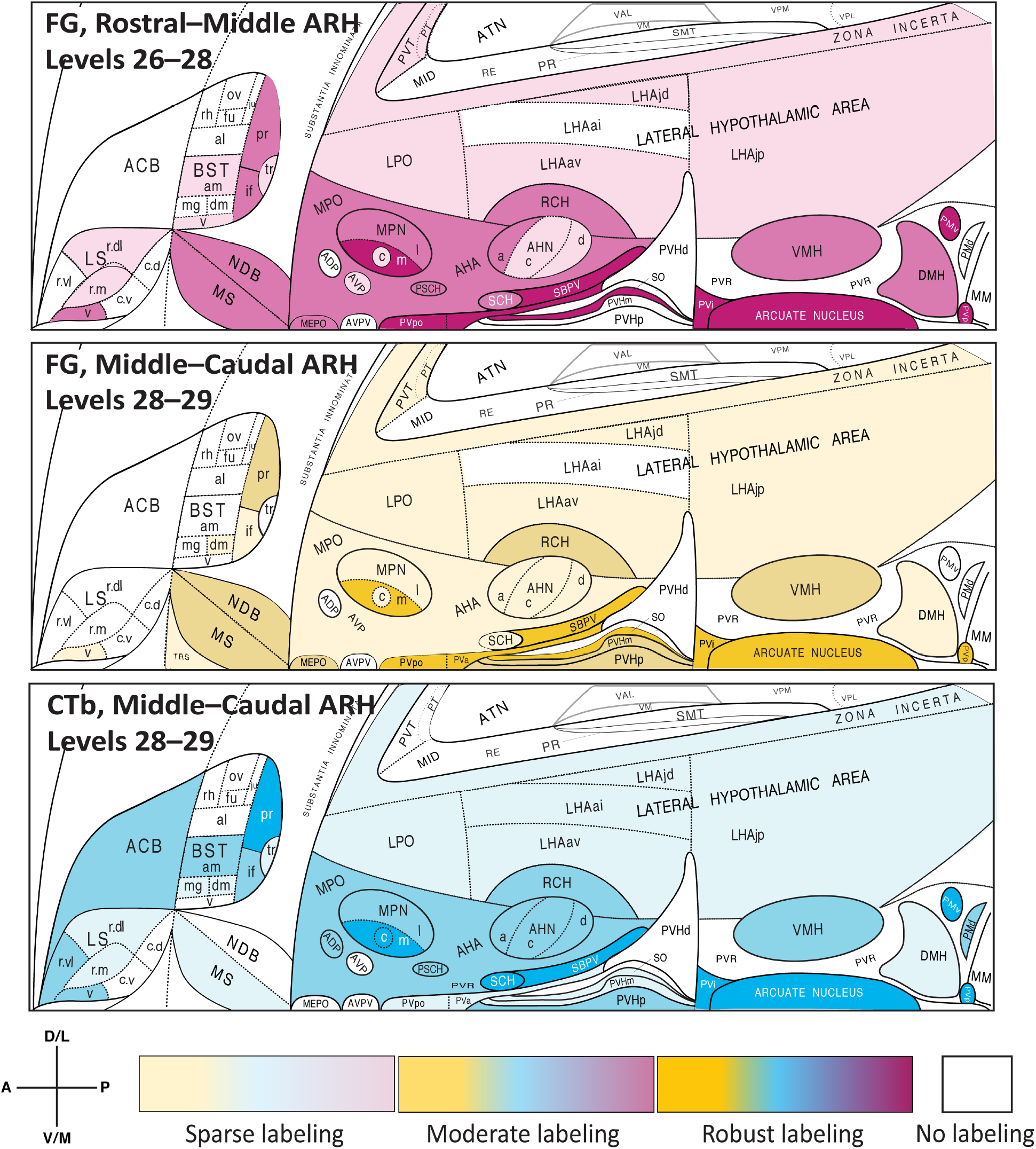
Provisional choropleth flatmaps of unpublished data (Martinez et al. 2015; 2016) are furnished here to illustrate differences in the distributions and densities of retrogradely labeled neurons traced from rostral versus caudal locations within the rat ARH using two different retrograde tracers across three test subjects (adult male Sprague-Dawley rats). The upper two panels show distributions and densities for neurons traced retrogradely from ARH injection sites containing deposits of the retrograde tracer, Fluorogold (FG). The bottom (*third from top*) panel is from a test subject receiving the retrogradely transportable tracer, cholera toxin subunit B (CtB). Note the locations within the ARH of the deposits, which were mapped to specific atlas levels of Swanson (2004) that contain the ARH. The deposit in the case shown in the upper panel was localized to the anterior half (rostral–middle) of the ARH, whereas the lower two panels were from subjects receiving deposits mapped to the posterior third (middle–caudal) of the ARH. Note that at atlas level 28 there was overlap of injection deposits for all three cases. Light to heavy shading for each panel corresponds to sparse to robust numbers of retrogradely labeled cells for each sub-region, with no shading indicating an absence of cells in that region. While many regions show similar densities and distributions of afferent input as would be expected from an overlapping series of injection deposits, there are also clear significant differences in such input to the rostral versus caudal portions of the ARH; most notably, the dorsomedial subdivision of the bed nuclei of the stria terminalis (BSTdm), the central part of the medial preoptic nucleus (MPOc), the periventricular part of the paraventricular hypothalamic nucleus (PVHp), and the dorsal premammillary nucleus (PMd). These differences in afferent distribution and density suggest that ARH recipient neurons are heterogeneous in at least certain functions (and therefore may differ in cellular state or phenotype), underscoring the need for accurate ARH sampling along the rostrocaudal axis of the nucleus for transcriptomic, proteomic or peptidomic studies. The flatmaps from Swanson (2004) (and available at https://larrywswanson.com) are reproduced and modified here under the conditions of a Creative Commons BY-NC 4.0 license (https://creativecommons.org/licenses/by-nc/4.0/legalcode).

## 8: Concluding remarks and future directions

In this article, we have surveyed the historical antecedents of high throughput technologies to extract molecular information from the brain, focusing on studies of the hypothalamus. After surveying selected articles reporting high throughput transcriptomic, proteomic and peptidomic studies of the hypothalamus or its sub-regions, we discussed the importance of LCM and digital atlasing methods in facilitating the anchoring (mapping) of such information to a tractable spatial model of the brain. In doing so, we build upon earlier efforts to link molecular information with spatial locations in the brain in a large-scale manner, such as grid-based mapping based on voxelation methods (Chin et al. 2007; C. Park et al. 2009; Petyuk et al. 2007; 2010) and analysis of gene expression patterns in the hypothalamus from the rich repository of *in situ* hybridization data within the Allen Brain Atlas (Olszewski et al. 2008).

### 8.1: Future directions in data management

We also apply the topics presented here to previous discussions we have raised concerning automated, informatics-based management of neuroscientific data; for example, using electronic laboratory notebooks to perform digital mapping and documentation of analyzed datasets (see Fig. 4 of Burns et al. 2003; Khan et al. 2006). As greater sophistication is brought to bear using methods that combine neuroanatomical tract tracing with molecular analysis of the traced projection neurons (e.g., see Pomeranz et al. 2017), both the mapping and management of data concerning such projection systems will become even more streamlined. An important aspect of developing informatics tools and methodologies is that much of the information used by expert biologists is technically specified, but informally defined. Naturally, expert neuroscientists are trained to understand the spatial structure of a published brain atlas, a flatmap, or a stained histological slide without requiring an explicitly defined logical representation. Informatics systems, however, must be grounded in a well-defined ontological model, which inevitably leads to some disagreement concerning the optimal design of such ontologies for neuroanatomical data. There are number of approaches put forward within the neuroinformatics community to represent mapped neuroanatomical data in ontologies. These include representations with a neuroimaging focus (Nichols et al. 2010); philosophically grounded approaches to neuroanatomy (Osumi-Sutherland et al. 2012); or comprehensive, cross-species methods for neuroanatomical phenotype (Haendel et al. 2014; Mungall et al. 2017; also see Deans et al. 2015). It is important to note that selecting an appropriate formalization can have a deep impact on how a neuroinformatics system functions, and we feel that any formalization used to represent the data described in this chapter should reflect the expertise and practices of experimental scientists working in this field. Thus, we recommend lightweight, data-centric formalizations that mirror scientists’ use of standard atlases, such as the Allen Brain Atlas portal (Sunkin et al. 2013). Neuroscientific knowledge carries a structured context that is inherited from the experimental design that ultimately generates the data. One methodology for representing this context in a general way is based on the relationships between independent and dependent variables within studies. This may serve as a convenient framework for describing neuroanatomically grounded data by treating the location of the phenomena of interest in the brain simply as one of several independent variables that describe the context of a particular datum (Russ et al. 2011; Tallis et al. 2011).

### 8.2: Future directions in imaging

Though it has not yet achieved the mesoscale resolution required to permit detailed mapping of most molecules, mass spectrometry imaging (MSI) – including imaging based on matrix assisted laser desorption technology (Karas et al. 1987), known as MALDI (Spengler et al. 1994; Caprioli et al. 1997) – will hopefully provide investigators the ability to rapidly sample the molecular landscape of the brain while simultaneously facilitating the preservation of the provenance of this molecular information at a resolution comparable to our proposed methods to map such information (for reviews on MALDI, see Cornett et al. 2007; Römpp et al. 2010; and Shariatgorji et al. 2014b). MALDI has now been performed for single neurons (Schwartz et al. 2003) and tissue sections (e.g., Heijs et al. 2015), including sections containing hypothalamus (Altelaar et al. 2006; Groseclose et al. 2007; I. Yao et al. 2008; Shariatgorji et al. 2014a) and pituitary (Altelaar et al. 2007). It is now being applied for metabolomics studies of the brain as well (Esteve et al. 2016). Modifications of the original method, including MALDI Fourier Transform Ion Cyclotron Resonance (MALDI FTICR; Spraggins et al. 2016), offer greater mass resolution and accuracy. A promising future direction for MALDI with respect to mapping of molecular information to canonical atlases is the recently reported strategy of combining MALDI with LCM and LC-MS/MS on the same brain section (Dilillo et al. 2017), which would facilitate the retention of provenance information for the molecular datasets mined from the section. Similarly, image fusion strategies that create one image of a tissue section from two registerable source images produced by two separate imaging modalities (MALDI, optical microcopy) also hold great promise for mapping molecular information (van de Plas et al. 2015). Other modalities, such as Raman spectroscopic imaging (Manciu et al. 2013), may offer additional opportunities for high spatial resolution analysis of molecular datasets in the brain.

### 8.3: Future directions in molecular analysis

Alongside developments in imaging technologies are enhanced technologies that allow for spatially resolved molecular sampling of tissue (see Crosetto et al. 2015, for a review). For example, fluorescent *in situ* sequencing (FISSEQ) of RNA has been developed for intact tissue samples (J. H. Lee et al. 2014; 2015). Similarly, Ståhl et al. (2016) have reported the novel use of arrayed reverse transcription primers accompanied by unique positional barcodes, which can be used to generate RNA-sequencing data directly on tissue slides in a manner that preserves the location of the information (also see Navarro et al. 2017). Additionally, single-cell transcriptomic analysis can be performed on individual nuclei obtained from fixed tissue; these nuclei are sorted after tissue dissociation procedures via fluorescence-activated cell sorting (FACS) or nucleic acid barcoding. For example, Lake et al. (2016) characterized the single nuclear transcriptomes of cerebral cortical neurons from fixed *post mortem* human brain. Habib et al. (2017) used barcoded beads to sort individual nuclei taken from fresh or frozen brain samples from mouse and human, and developed a microfluidic device that enables the sorting process.

A key future direction would be to integrate spatially resolved transcriptomics procedures and single-cell sequencing efforts into a pipeline that allows for the retention and mapping of the locations from where the samples originate with respect to canonical brain atlases. Along these lines, the Retro-TRAP technology developed by Jeffrey Friedman and colleagues (Ekstrand et al. 2014; Nectow et al. 2015; Pomeranz et al. 2017; derived from the original TRAP technology to identify activated neurons: Knight et al. 2012) to retrogradely label neurons with GFP constructs and then capture translating mRNAs from these neurons using anti-GFP nanobodies (i.e., single-domain antibodies), could potentially allow for projections being mapped for neurons from which single-cell molecular information can also be harvested in a spatially documentable manner. A current limitation of the method for such purposes is that fresh not frozen tissue needs to be harvested to generate sufficient RNA yields, precluding the freezing of tissue sections in preparation for LCM.

### 8.4: Future directions in mapping

A larger issue concerning the mapping of molecular information is the need to change the scientific culture so that best practices of reporting molecular information in the brain include procedures to map the information to standardized atlases. At present, this is not a common practice by most investigators in neuroscience. Such changes in culture would greatly accelerate the integration of datasets among researchers, and the need to do so is now more critical than ever, given the deluge of spatial, molecular data that has already been shared in repositories such as the Allen Brain Atlas (http://www.brain-map.org) and the GENSAT Project (http://www.gensat.org) (also see Gray et al. 2004; Magdaleno et al. 2006; Lein et al. 2007; Shimogori et al. 2010).

### 8.5: Final remarks

For all of these and future advancements, it will remain critical to preserve information about the native lands from which so many molecules become expatriated, lest the information provided by these molecular datasets fails to link up with the larger neuronal information networks from which they came. Mapping the sampled tissue will provide the critical information that will bridge the gap between the systems biology of molecular information networks on the one hand, and the systems neuroscience of cellular information networks on the other. Without such a bridge, these domains of inquiry may never converge to form a unifying model of a dynamic brain, replete with diverse molecular citizens hailing from different but interconnecting cells, and communicating across local and regional boundaries to signal their neighbors, both near and far.

## Funding

Work in the UTEP Systems Neuroscience Laboratory is supported by grants awarded to AMK from the National Institutes of Health (NIH; SC3GM109817; SC1GM127251), the Howard Hughes Medical Institute (UTEP PERSIST; PI: S. Aley), and the UTEP Office of Research and Sponsored Projects (Grand Challenges Award). This work is also supported by funds awarded to the Border Biomedical Research Center by the National Institute of Minority Health and Health Disparities of the NIH (5G12MD007592). AHG is supported by the Research Initiative for Scientific Enhancement (RISE) Graduate Fellowship program of the NIH (R25GM069621). AM has been supported by UTEP PERSIST funds and an NSF GK–12 fellowship. Some data in this study were also based upon work supported by the Office of Research and Development, Medical Research Service, Department of Veterans Affairs (VA); specifically, by Merit Review Awards 1l01BX001213-01A1 and BX004102-01 from the United States (U.S.) Department of Veterans Affairs Biomedical Laboratory Research and Development Service to JEB as well as NIH R01DK115976 to JEB. The contents do not represent the views of the U.S. Department of Veterans Affairs or the United States Government. This study was also supported by the University of Washington Diabetes Research Center Cellular and Molecular Imaging Core, which is supported by NIH grant P30DK017047. The contribution by GAPCB to this work was funded by the Defense Advanced Research Projects Agency (DARPA) Big Mechanism program under Army Research Office (ARO) contract W911NF-1-0436 and by NIH grant R01LM012592.

## Acknowledgments

We thank Dr. Sabiha Khan (UTEP) for thoughtful discussion on the organization of the manuscript, and Dr. Harold Gainer (National Institute of Neurological Disorders and Stroke) for his timely feedback. We also acknowledge our debt to the late Dr. Claude F. Baxter, who served as Emeritus Professor of Psychiatry and Biobehavioral Sciences at the UCLA Brain Research Institute and past historian of the American Society for Neurochemistry, for having generously provided AMK access to his personal library of seminal works in neurochemistry. His kindness and hospitality are treasured memories. We would also like to acknowledge the contributions of Dr. Rebecca Hull and Nishi Gill for the images provided in Figures 2B and 2C. Finally, we thank Dr. Alexander C. Jackson (University of Connecticut) for providing us with access to unpublished data from his single-cell transcriptomic studies of neuron populations in the mouse lateral hypothalamic area. This article is dedicated to the memory of Dr. John H. Ashe (University of California at Riverside), whose instruction and mentorship have deeply informed this narrative.

## Abbreviations used

ACB: nucleus accumbens
AchE: acetylcholinesterase
ADP: anterodorsal preoptic nucleus
AgRP: Agouti-Related Peptide
AHN: anterior hypothalamic nucleus
AHNa: anterior hypothalamic nucleus, anterior part
AHNc: anterior hypothalamic nucleus, central part
AHNd: anterior hypothalamic nucleus, dorsal part
AHNp: anterior hypothalamic nucleus, posterior part
AP: area postrema
ARH: arcuate hypothalamic nucleus
ATN: anterior nuclei, dorsal thalamus
AVP: anteroventral preoptic nucleus
AVPV: anteroventral periventricular nucleus hypothalamus
BST: bed nuclei of the stria terminalis
BSTal: bed nuclei of the stria terminalis, anterior division, anterolateral area
BSTam: bed nuclei of the stria terminalis, anterior division, anteromedial area
BSTdm: bed nuclei of the stria terminalis, anterior division, dorsomedial nucleus
BSTfu: bed nuclei of the stria terminalis, anterior division, fusiform nucleus
BSTif: bed nuclei of the stria terminalis, posterior division, interfascicular nucleus
BSTju: bed nuclei of the stria terminalis, anterior division, juxtacapsular nucleus
BSTmg: bed nuclei of the stria terminalis, anterior division, magnocellular nucleus
BSTov: bed nuclei of the stria terminalis, anterior division, oval nucleus
BSTpr: bed nuclei of the stria terminalis, posterior division, principal nucleus
BSTrh: bed nuclei of the stria terminalis, anterior division, rhomboid nucleus
BSTtr: bed nuclei of the stria terminalis, posterior division, transverse nucleus
BSTv: bed nuclei of the stria terminalis, anterior division, ventral nucleus
C.a.: anterior commissure
CCK1R: cholecystokinin 1 receptor
C.f.d.: fornix
Ch. Opt.: optic chiasm
CRH: corticotropin-releasing hormone
CTB: cholera toxin subunit b
DMH: dorsomedial hypothalamic nucleus
EGFP: enhanced green fluorescent protein
FG: Fluorogold
fx: fornix
GFP: green fluorescent protein
HNS: hypothalamo-neurophypohysial system
I: internuclear area, hypothalamic periventricular region
KO: knockout
LCM: laser capture microdissection
LHA: lateral hypothalamic area
LHAai: lateral hypothalamic area, anterior region, intermediate zone
LHAav: lateral hypothalamic area, anterior region, ventral zone
LHAd: lateral hypothalamic area
LHAjd: lateral hypothalamic area, juxtadorsomedial region
LHAjp: lateral hypothalamic area, juxtaparaventricular region
LHAjvd: lateral hypothalamic area, juxtaventromedial region, dorsal zone
LHAjvv: lateral hypothalamic area, juxtaventromedial region, ventral zone
LHApc: lateral hypothalamic area, parvicellular region
LHAsfa: lateral hypothalamic area, subfornical region, anterior zone
LPO: lateral preoptic area
LS: lateral septal nucleus [Cajal]
LSc.d: lateral septal nucleus, caudal part, dorsal zone
LSc.v: lateral septal nucleus, caudal part, ventral zone
LSr.dl: lateral septal nucleus, rostral part, dorsolateral zone
LSr.m: lateral septal nucleus, caudal part, medial zone
LSr.vl: lateral septal nucleus, rostral part, ventrolateral zone
LSv: lateral septal nucleus, ventral part [Risold-Swanson]
MC4-R: melanocortin 4 receptor
ME: median eminence
MEex: median eminence, external lamina
MEin: median eminence, internal lamina
MEPO: median preoptic nucleus
MID: midline nuclei, dorsal thalamus
MM: medial mammillary nucleus, body
MNs: magnocellular neurons
MPN: medial preoptic nucleus
MPNc: medial preoptic nucleus, central part
MPNl: medial preoptic nucleus, lateral part
MPNm: medial preoptic nucleus, medial part
MPO: medial preoptic area
MS: medial septal nucleus [Cajal]
μ-array: microarray
NDB: diagonal band nucleus [Broca]
NPY: neuropeptide Y
NTS: nucleus of the solitary tract
opt: optic tract
OT: oxytocin
PCR: polymerase chain reaction
PFA: paraformaldehyde
PMd: dorsal premammillary nucleus
PMv: ventral premammillary nucleus
POMC: pro-opiomelanocortin
PR: perireuniens nucleus
PSCH: suprachiasmatic preoptic nucleus
PT: paratenial nucleus
PVH: paraventricular hypothalamic nucleus
PVHd: paraventricular hypothalamic nucleus, descending division
PVHf: paraventricular hypothalamic nucleus, descending division, forniceal part
PVHm: paraventricular hypothalamic nucleus, magnocellular division
PVHmpd: paraventricular hypothalamic nucleus, medial parvicellular part, dorsal zone
PVHp: paraventricular hypothalamic nucleus, parvicellular division
PVHpv: paraventricular hypothalamic nucleus, periventricular part
PVi: periventricular hypothalamic nucleus, intermediate part
PVp: periventricular hypothalamic nucleus, posterior part
PVpo: preoptic periventricular nucleus
PVT: paraventricular thalamic nucleus
PVR: hypothalamic periventricular region
qPCR: quantitative polymerase chain reaction
RCH: retrochiasmatic area, lateral hypothalamic area
RE: nucleus reuniens [Malone]
REcd: nucleus reuniens, caudal division, dorsal part
REcm: nucleus reuniens, caudal division, medial part [Gurdjian]
REcp: nucleus reuniens, caudal division, posterior part
RIN: RNA integrity number
SBPV: subparaventricular zone hypothalamus
SCH: suprachiasmatic nucleus [Spiegel-Zwieg]
SFO: subfornical organ
SMT: submedial nucleus thalamus
SO: supraoptic hypothalamic nucleus
SOr: supraoptic nucleus, retrochiasmatic part
S.t.: infundibular stalk
sup: supraoptic commissures
TH: tyrosine hydroxylase
T.M.: tractus Meynert (fasciculus retroflexus)
TUi: tuberal nucleus, intermediate part
TUsv: tuberal nucleus, subventricular part
V3h: third ventricle, hypothalamic part
V.d’A.: tract of Vicq D’Azyr (mammillothalamic tract)
vlt: ventrolateral hypothalamic tract
VMH: ventromedial hypothalamic nucleus
VMHa: ventromedial hypothalamic nucleus, anterior part
VMHc: ventromedial hypothalamic nucleus, central part
VMHdm: ventromedial hypothalamic nucleus, dorsomedial part
VMHvl: ventromedial hypothalamic nucleus, ventrolateral part
VP: vasopressin
VPL: ventral posterolateral nucleus thalamus, principal part
VPM: ventral posteromedial nucleus thalamus, principal part

